# Meanders as a scaling motif for understanding of floodplain soil microbiome and biogeochemical potential at the watershed scale

**DOI:** 10.1101/2020.05.14.086363

**Authors:** Paula B. Matheus Carnevali, Adi Lavy, Alex D. Thomas, Alexander Crits-Christoph, Spencer Diamond, Raphaeël Meéheust, Matthew R. Olm, Allison Sharrar, Shufei Lei, Wenming Dong, Nicola Falco, Nicholas Bouskill, Michelle Newcomer, Peter Nico, Haruko Wainwright, Dipankar Dwivedi, Kenneth H. Williams, Susan Hubbard, Jillian F. Banfield

## Abstract

Biogeochemical exports of C, N, S and H_2_ from watersheds are modulated by the activity of microorganisms that function over micron scales. This disparity of scales presents a substantial challenge for development of predictive models describing watershed function. Here, we tested the hypothesis that meander-bound regions exhibit patterns of microbial metabolic potential that are broadly predictive of biogeochemical processes in floodplain soils along a river corridor. We intensively sampled floodplain soils located in the upper, middle, and lower reaches of the East River in Colorado and reconstructed 248 draft quality genomes representative at a sub-species level. Approximately one third of the representative genomes were detected across all three locations with similar levels of abundance, and despite the very high microbial diversity and complexity of the soils, ~15% of species were detected in two consecutive years. A core floodplain microbiome was enriched in bacterial capacities for aerobic respiration, aerobic CO oxidation, and thiosulfate oxidation with the formation of elemental sulfur. We did not detect systematic patterns of gene abundance based on sampling position relative to the river. However, at the watershed scale meander-bound floodplains appear to serve as scaling motifs that predict aggregate capacities for biogeochemical transformations in floodplain soils. Given this, we conducted a transcriptomic analysis of the middle site. Overall, the most highly transcribed genes were *amoCAB* and *nxrAB* (for nitrification) followed by genes involved in methanol and formate oxidation, and nitrogen and CO_2_ fixation. Low soil organic carbon correlated with high activity of genes involved in methanol, formate, sulfide, hydrogen, and ammonia oxidation, nitrite oxidoreduction, and nitrate and nitrite reduction. Thus, widely represented genetic capacities did not predict *in situ* activity at one time point, but rather they define a reservoir of biogeochemical potential available as conditions change.

## Introduction

Watersheds are geographic areas that capture precipitation that is ultimately discharged into rivers and other larger water bodies. Of particular interest are watersheds in mountainous regions, as these are major sources of freshwater ^1, 2^. Within mountainous watersheds, complex interactions among vegetation, hydrology, geochemistry, and geology occur within and across watershed compartments, including across bedrock-soil-vegetation compartments of terrestrial hillslopes, across terrestrial-aquatic interfaces and within the fluvial system itself. Interactions within a reactive watershed typically vary as a function of disturbance as well as landscape position and topography. For example, interactions in an alpine region of a mountainous watershed are likely to be quite different from a lower montane floodplain region ^3^. Floodplains, which extend from the river banks to the base of hillslopes, comprise the riparian zone (a vegetated interface between the river channel and the rest of the ecosystem), and are notable as they integrate inputs from all watershed compartments. They also display depositional gradients and features associated with past and current river channel positions. Unlike hillslopes, floodplains receive water and constituents either by surface runoff or groundwater discharge. They are typically significantly impacted by changes in river conditions and can be inundated when river flow and stage increases following snowmelt. Consequently, floodplains are dynamic compartments in which hydrobiogeochemical processes vary seasonally and potentially spatially. Overall, floodplains are important watershed regions in which microbial activity can modulate the form and abundance of nutrients and contaminants derived from hillslopes and river water prior to their export from the watershed.

Here, we conducted a study of floodplain soils of the mountainous East River (CO) watershed to investigate how patterns in the distribution of soil microorganisms and their associated functions and activities can induce geochemical gradients that impact riverine nutrient and contaminant fluxes. We tested a ‘system-of systems’ approach ^4^ wherein meander-bound regions were selected as scaling motifs (repeating patterns along the river that can be used for ecosystem modeling at the watershed scale), in which microbially-mediated biogeochemical processes that are shaped by reactions occurring at the micron-scale might be representative of processes throughout the floodplain. Detailed analyses of meander-bound floodplain soils may reveal patterns that approximate watershed processes at the tens of kilometers scale, and could provide much needed input for watershed hydrobiogeochemical models. This study applied genome-resolved metagenomic and metatranscriptomic bioinformatics methods to large nucleic acid sequence datasets to investigate microbial community composition and distribution and to infer capacities for microbially-mediated C, S, H and N cycling and *in situ* activity in floodplain soil microbial communities.

## Results

### Metagenomes overview

Three meander-bound floodplains following the meandering pattern of the East River (**Fig. 1a**) were chosen for this study: one upstream (meander-bound floodplain G (Floodplain G); **Fig. 1b**), one midstream (meander-bound floodplain L (Floodplain L); **Fig. 1c**), and one downstream (meander-bound floodplain Z (Floodplain Z); **Fig. 1d**). Sample number was prioritized over sequencing depth to better resolve the types and distribution patterns of the most abundant organisms across the meander-bound floodplains (‘floodplains’ subsequently). An average 6.4 giga base pairs (Gbp; 3.2 – 11.5 Gbp) of sequencing data was obtained from 90 DNA extractions out of 94 floodplain soil samples collected in 2015. An average 12 Gbp (6.0 – 15.0 Gbp) of sequencing data was obtained from the other four samples. In total, ~0.6 Tbp of DNA sequence information was acquired from samples collected in 2015. Our strategy aimed to capture the most abundant members of the microbial community (instead of the whole community), so it is not surprising that an average 13% (3 - 30%) of reads mapped to their respective assemblies (Supplementary Table 1). This result also reflects the large (but variable) tail on the abundance distribution of microorganisms in soil ^5^; and the communities captured by the smaller assemblies resulting from samples with lower sequencing depth.

**Figure 1.**
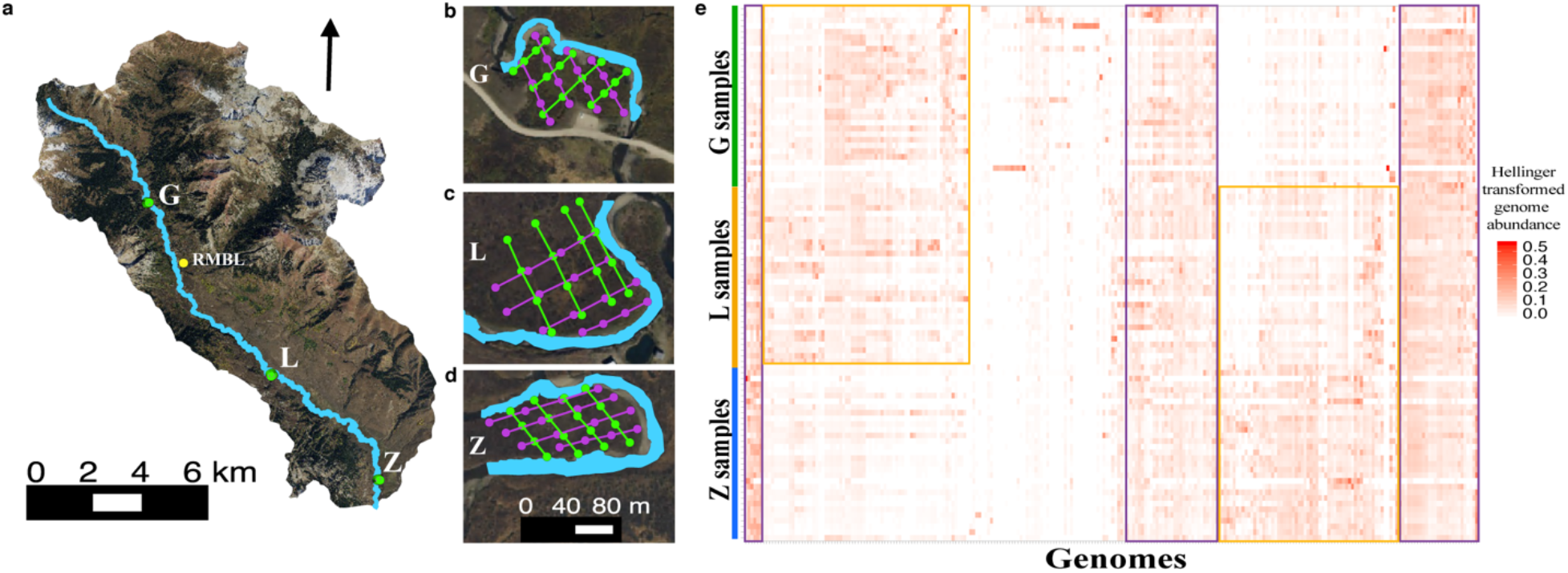
(**a**) Overview of the East River, CO study site, highlighting the three sampled floodplains (green dots) and the Rocky Mountain Biological Laboratory (RMBL, yellow dot), (**b**) meander-bound floodplain G, (**c**) meander-bound floodplain L, (**d**) meander-bound floodplain Z. Sampling sites as green and purple dots along two sets of four transects. One set of transects in one direction (in green), and the second set of transects along another direction (in purple). (**e**) Hellinger transformed abundance of dereplicated genomes across samples based on cross-mapping. Genomes and samples clustered by average linkage and Euclidean distance respectively.

We constructed 1,704 draft genomes from three floodplain datasets. About one third (622) of these genomes were classified as draft quality (237 from floodplain G, 150 from floodplain L, and 235 from floodplain Z). After dereplication at 99% average nucleotide identity (ANI) within each floodplain and correction of local assembly errors, we recovered 375 distinct genomes (173 from G, 94 from L and 108 from Z). Dereplication across floodplains at 98% ANI generated a final set of 248 representative genomes for further analyses, predominantly ≥ 70% complete (Supplementary Table 2) and 46% of which were near-complete (≥ 90%).

We assessed our genome recovery effectiveness by comparing the number of genomes recovered to a secondary metric for quantifying unique species, the number of unique ribosomal protein L6 (rpL6) marker sequences within unbinned assemblies (see **Methods**). The marker rpL6 has been shown to have high recoverability and species delineation accuracy ^6^, relative to methods such as full genome ANI. From the 94 metagenomes, we detected 930 distinct organisms based on rpL6 sequences clustered at 97.5% nucleotide identity. However, 571 of the distinct rpL6 sequences were on fragments with coverage that is too low for comprehensive genome sampling (<7 X coverage given our sequencing depth). The disparity relative to 248 reconstructed representative genomes relative to 359 rpL6 on contigs at >7 X coverage is attributed to significant challenges associated with genome recovery from soil.

Candidate draft genomes were generated for almost all the organisms present at > 5-10 X coverage in each sample. However, on average, only 5.5% of the total read dataset was stringently mapped (2 mismatches per read of the pair) to the 248 genomes. This is not surprising, given that most sequencing allocations per sample were sufficient to genomically sample only organisms at > ~0.25% relative abundance, and the most abundant organisms in each sample comprised only a few percent of the community.

In 2016, we returned to one of the floodplain sites (floodplain L) to collect samples for metatranscriptomics. We performed additional genomic sequencing from 19 of the original 32 sites (see **Methods;** Supplementary Table 1) to provide a reference database for transcript mapping. These new DNA samples were sequenced at an average 3.7 Gbp per sample (2.7 – 4.7 Gbp), for a total ~ 0.2 Tbp of sequencing. The RNA samples were sequenced at an average 10.8 Gbp per sample (3.2 - 15.1 Gbp) for a total of ~ 0.15 Tbp of sequencing. A total of 299 draft genomes were recovered from these samples, 123 of which passed our quality thresholds after curation. To examine stability across time we pooled the 2015 and 2016 genome sets and dereplicated (at 95% ANI) the combined set of 371 genomes, generating 215 genomes representative of distinct species. Notably, 32 species-level groups were detected in both years and 29 were only detected in 2016 (Supplementary Table 3).

### Distribution of organisms within and across meander-bound floodplains

To assess the presence of a representative genome in a sample we relied on the sensitivity of read mapping to the dereplicated genome set. Based on our threshold for detection, about one-third of the genomes were from organisms that were consistently found across floodplains at similar levels of abundance (Purple boxes in **Fig. 1e**). Regardless of their level of abundance, or which floodplain a representative genome was reconstructed from, the genomes were present in a median > 75% of the samples (78% of upstream floodplain G samples, 84% of mid-stream floodplain L samples, and 87% of downstream floodplain Z samples; Supplementary Figure 1a). Additionally, the 248 organisms were present in the majority (median 88-91%) of the other samples from the same floodplain from which the genome was reconstructed from (Supplementary Figure 1b).

Except for some genomes reconstructed from two floodplain G samples, the rest of the genomes were from organisms that shared more similar abundance levels if the floodplains were closer together within the river corridor (Yellow boxes in **Fig. 1e**). More specifically, floodplains G and L or floodplains L and Z shared more organisms than floodplains G and Z, which are located in the upper and lower reaches respectively. Additionally, floodplain G is narrow, and is at times completely flooded, floodplain L is wider and may only flood partially, whereas floodplain Z is the widest and least prone to flooding.

Finally, we examined the number of samples where members of a 98% ANI genome cluster were reconstructed from. The 248 genome clusters contained genomes reconstructed from between 2 to 39 samples. The largest genome set was for a large group of Betaproteobacteria strains generally related to strains detected in other environments such as soil, sediment and water (Supplementary Figure 2; Supplementary Data 1). Genomes were reconstructed from two thirds of all samples from floodplain L. This result indicates that strains belonging to this Betaproteobacterial clade may play important roles in floodplain biogeochemistry (**Fig. 2**), especially in soils associated with floodplain L.

**Figure 2.**
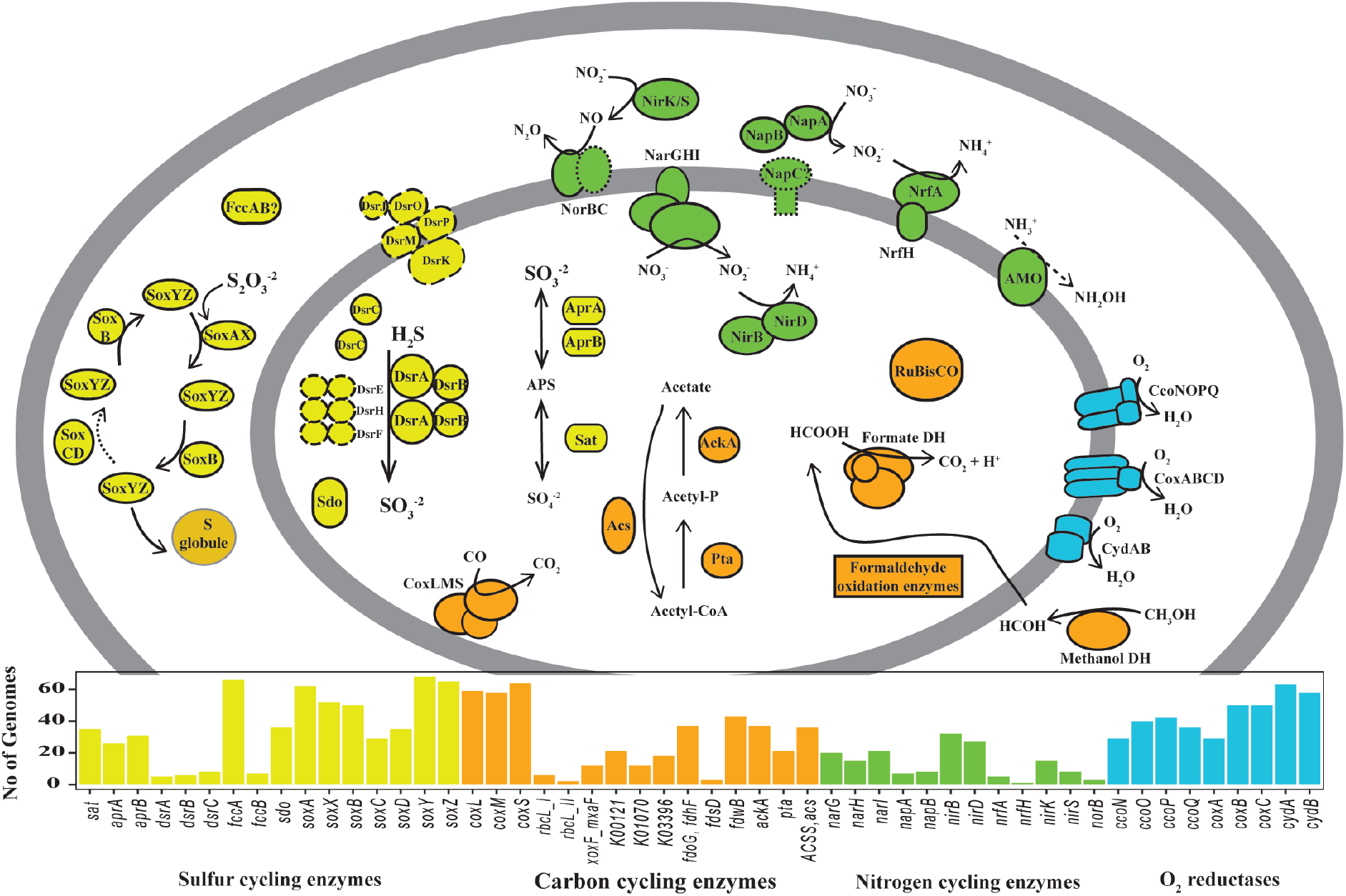
Diagram depicting Betaproteobacteria genomes and environmentally relevant capacities encoded by representatives of 98% ANI clusters. Note that no single genome harbors all of these genes, but combinations of them instead (Supplementary Table 4). Some genomes harbor methanol dehydrogenases that are potentially able to turn methanol directly into formate (XoxF type). Enzymes delineated with solid lines were predicted using KOFAM HMMs, and the number of genomes (> 1*) encoding those genes are shown in the bars plot. Enzymes that were predicted using methods as part of ggKbase are shown with dashed lines (long dashes) and enzymes or subunits that are presumably encoded are shown with dotted lines. For more information about metabolic potential see **Methods**. *AMO was included in this diagram even though it was detected in only 1 genome, to indicate aerobic ammonia oxidation is also possibly carried out by members of this clade.

### Taxonomic composition of the community

Based on the 248 representative genomes detected in each sample, the phylum- or class-level community composition was broadly consistent both within and across floodplains (**Fig. 3a**). This is also reflected in measures of alpha diversity between sites, as there was no significant difference in Shannon’s diversity indices or unique number of organisms (Supplementary Figure 3). Some exceptions to this were Candidate Phyla Radiation (CPR) bacteria that seemed to be detected mostly in floodplain Z, while Thaumarchaeota seemed least present in this floodplain. Additionally, the number of the 248 genomes detected in each sample varied from sample to sample (min = 35 and max = 212). We detected particularly low numbers of genomes in five samples (T157 and T800 from floodplain G and T133, T266 and T620 from floodplain Z), although only samples T133 and T266 from floodplain Z may have been affected by lower sequencing depths (Supplementary Table 1).

**Figure 3.**
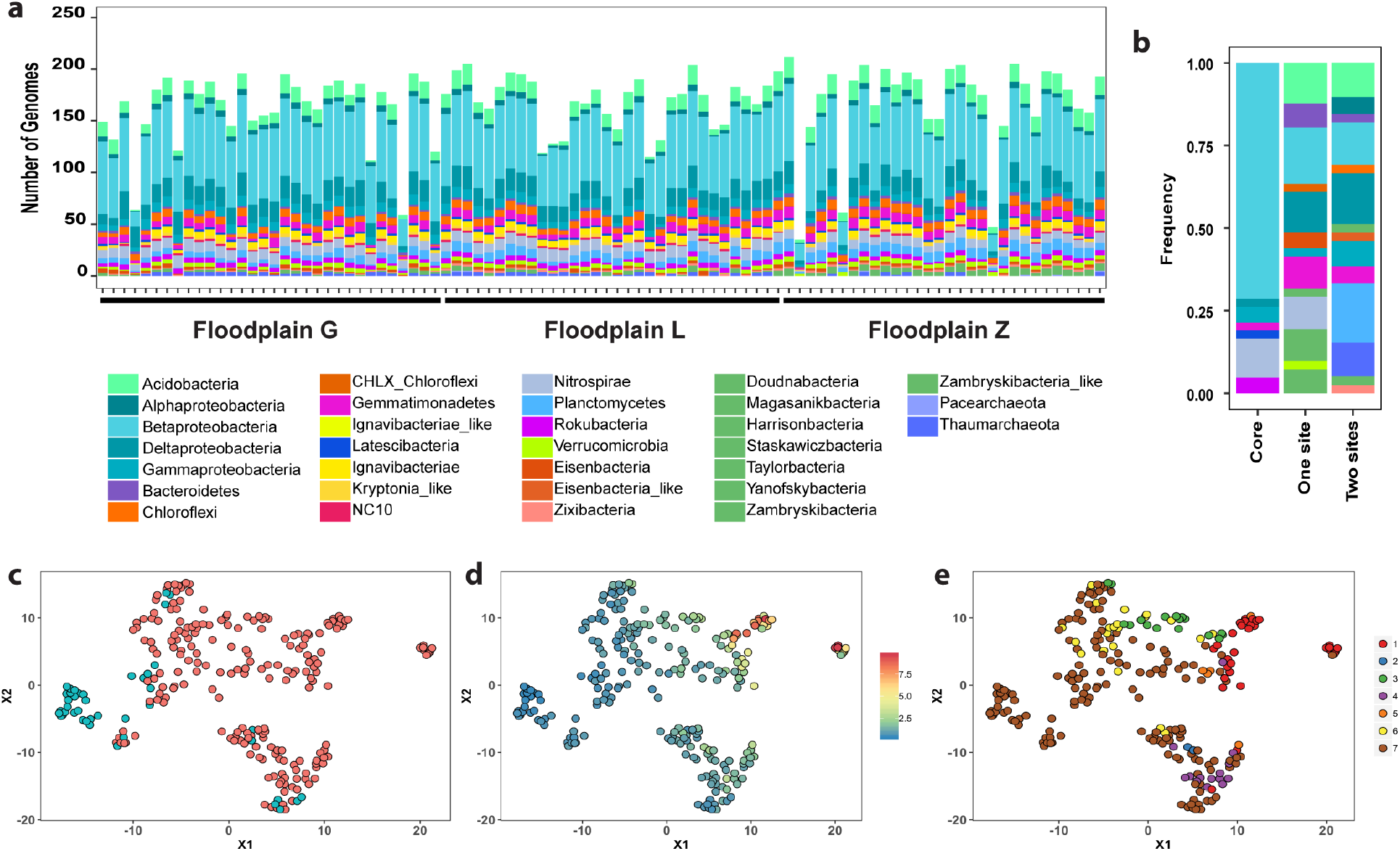
**(a)** Taxa at the phylum or class level detected across samples and floodplains, samples are in numerical order (Supplementary Table 1) from the upstream to the downstream floodplain. **(b)** Taxonomic composition of genomes in the core floodplain microbiome (core), genomes associated with 1 floodplain (one site), and genomes associated with two floodplains (two sites). UMAP showing clustering of Hellinger transformed genome abundances of **(c)** genomes in the core floodplain microbiome (n = 42; in teal) and genomes not in the core floodplain microbiome (n = 242; red), (**d**) and overlay of the coefficient of variation (ratio of standard deviation to the mean) of genome abundances across samples, and **(e)** overlay of genomes associated with individual, pairs, or all floodplains based on an Indicator Species Analysis (ISA). Genomes that were present in 89 samples or more (teal) were not associated with any particular floodplain by ISA (group 7 in brown) and their abundance displayed a low coefficient of variation across samples. ISA genome associations: with floodplain G (1), with floodplain L (2), with floodplain Z (3), with both floodplain G and floodplain L (4), with both floodplain G and floodplain Z (5), with both floodplain L and floodplain Z (6), not associated with any particular floodplain (7).

Betaproteobacteria was the group with the highest number of representative genomes (80) in all three floodplains. Other abundant taxa across floodplains included Deltaproteobacteria (27 representatives), Acidobacteria (21 representatives), Nitrospirae and Planctomycetes (both with 13 representatives), Gemmatimonadetes, Gammaproteobacteria, Chloroflexi and Ignavibacteria (12, 11, 11, and 10 respectively).

### Uncovering a core floodplain microbiome

To define a set of organisms representing a core floodplain microbiome we identified organisms that were detected in most sampled sites (≥ 89 of the 94 samples; 90t^h^ percentile), and whose abundance did not indicate a statistically significant enrichment in any specific floodplain (by Indicator Species Analysis (ISA); see **Methods** and Supplementary Table 5). This operational definition resulted in the identification of 42 high prevalence organisms with low variance abundance profiles across all 3 meander-bound sites, which we refer to as the core floodplain microbiome (**Fig. 3c**). In general, genomes with a low coefficient of variation of their abundance (blue dots in **Fig. 3d**) overlapped with genomes that did not display a statistically significant association with any given floodplain (group 7, brown dots in **Fig. 3e**), suggesting a wide distribution of these organisms across floodplains at similar abundance levels. The core floodplain microbiome was dominated by Betaproteobacteria, with lower abundances of Nitrospirae, Rokubacteria, Gemmatimonadetes, Gammaproteobacteria, Deltaproteobacteria, and Candidatus Letescibacteria (**Fig. 3b**).

Genomes from organisms not considered to be part of the core floodplain microbiome were associated with one floodplain (G, L, or Z; n = 41) or two floodplains (n = 39; **Fig. 3e**). Other genomes were not classified as part of the core microbiome because although they were not statistically associated with one or two floodplains, they were not detected in ≥ 89 samples (n = 126). Genomes affiliated with Acidobacteria, Bacteroidetes, and Chloroflexi were not part of the core floodplain microbiome and were associated with one or two floodplains. The ISA analysis supports the association of some CPR with one floodplain (*i.e.,* between floodplain Z and Yanofskybacteria, Taylorbacteria, Harrisonbacteria, Staskawiczbacteria, and Zambryskibacteria-like bacteria). Similarly, bacteria in the Verrucomicrobia were associated with one floodplain (Z). Alphaproteobacteria, Thaumarcheota, Planctomycetes, other CPR (*e.g.*, Zambryskibacteria and Doudnabacteria) and Eisenbacteria-like bacteria were associated with two floodplains.

### Geochemical functions, including those enriched in the core floodplain microbiome

To determine what role floodplain soil Bacteria and Archaea may play in nutrient exports to the East River, we investigated a set of pathways involved in biogeochemical cycling and the microorganisms potentially responsible for them. The biogeochemical processes investigated include oxidation/reduction reactions associated with nitrogen, sulfur and hydrogen, C1 compound metabolism (*e.g.,* CO_2_-fixation, CO oxidation, methanogenesis, methane oxidation, methanol oxidation, formate oxidation, methylamine oxidation, formaldehyde oxidation), H_2_ consumption or production, and the ability to use O_2_ as a terminal electron acceptor for aerobic respiration.

A set of HMMs was used to annotate genes encoding for individual protein subunits that make up key enzymes and complete or partial metabolic pathways. For a given ‘function’ (defined as the capacity to carry out a given biogeochemical transformation) to be encoded in a genome, certain criteria for presence had to be met (see **Methods**; Supplementary Table 4). A total of 32 functions comprised the final set of biogeochemical transformations under investigation (Supplementary Table 6). It is important to note that in some cases we also examined individual steps that are involved in a function, recognizing that some functions could be absent in a single genome because the pathway is carried out by multiple taxa (*i.e.,* steps are encoded in multiple genomes). For example, denitrification occurs in separate steps involving different enzymes, and these steps can be performed by multiple different organisms. Complete ammonia oxidation, anaerobic ammonia oxidation, and methanogenesis (of any kind), were not detected in the dereplicated genome set, although some intermediary steps may still be ecologically relevant. Therefore, some steps involved in these pathways were included in downstream analyses.

To study the distribution of the functions of interest among genomes and across floodplains, we determined whether a function was present or absent in each genome in addition to where genomes were detected within and across floodplain samples. To describe the distribution of functions, we calculated the proportion of genomes with a given function compared to the total number of genomes detected in a sample. We found that the ability to use oxygen as an electron acceptor (aerobic respiration) was the most prevalent function among genomes (a median of 70 - 85% of genomes in each sample), followed by acetate metabolism (a median of 40 - 65% of genomes in each sample), aerobic carbon monoxide (or other small molecule) oxidation, formate oxidation, and sulfide oxidation (a median of 30 - 50% of genomes in each sample; **Fig. 4a**). This set of functions was consistently present across all three floodplains, whether encoded by the same or different taxa.

**Figure 4.**
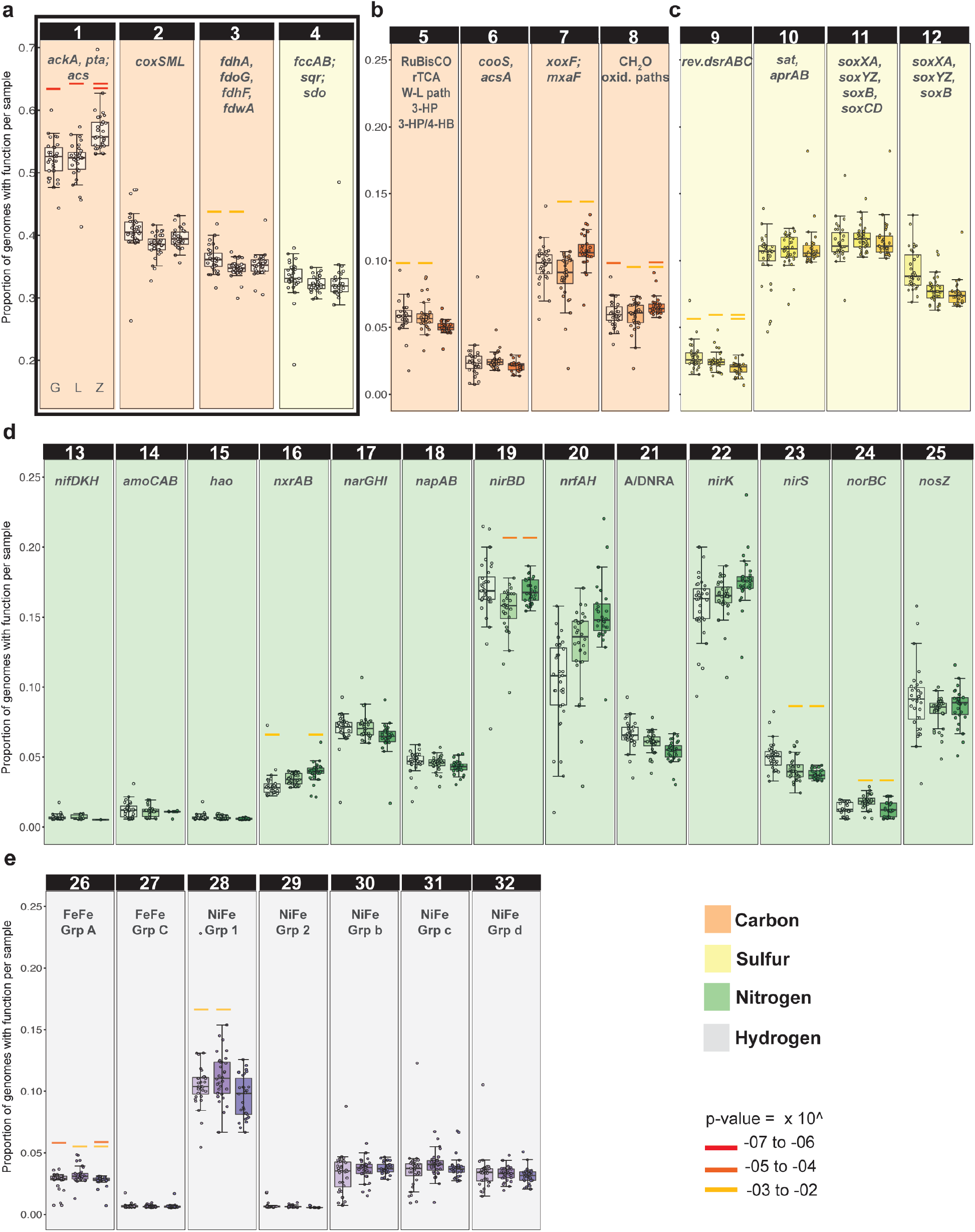
Proportion of representative genomes at the sub-species level with a function among genomes detected in each sample within each floodplain. In each panel the box plot on the left shows floodplain G, the box plot in the middle shows floodplain L, and the boxplot on the right shows floodplain Z. (**a**) Most abundant functions: **1**. Acetate formation, **2**. Oxidation of CO and other small molecules, **3**. Formate oxidation: CH_2_O_2_ to CO_2_ + H_2_, **4**. Sulfide oxidation: H_2_S to S^0^. (**b**) Geochemical transformations in the Carbon cycle: **5**. CO_2_ fixation pathways, **6**. Anaerobic CO oxidation, **7**. Methanol oxidation, **8**. Formaldehyde oxidation pathways (see Supplementary Table 4). (**c**) Geochemical transformations in the sulfur cycle: **9**. Sulfide oxidation (reverse *dsr*) from hydrogen sulfide: H_2_S to SO_3_^2−^, **10**. Sulfite oxidation to sulfate (or vice versa): SO_3_^2−^ to SO_4_^2−^, **11**. Thiosulfate oxidation without sulfur deposition: S_2_O_3_^2−^ to SO_4_^2^, **12**. Thiosulfate oxidation with sulfur deposition: S_2_O_3_^2−^ to SO_4_^2−^ + S^0^. (**d**) The nitrogen cycle: **13**. Nitrogen fixation: N_2_ to NH_3_, **14**. Ammonia oxidation: NH_3_ to NH_2_OH, **15**. Hydroxylamine oxidation (requires additional, undetermined enzyme): NH_2_OH to NO_2_^−^, **16**. Nitrite oxidation: NO_2_^−^ to NO_3_^−^ (reversible), **17**. Nitrate reduction (cytoplasmic): NO_3_^−^ to NO_2_^−^, **18**. Nitrate reduction (periplasmic): NO_3_^−^ to NO_2_^−^, **19**. Assimilatory nitrite reduction: NO_2_^−^ to NH_4_, **20**. Dissimilatory nitrite reduction: NO_2_^−^ to NH_4_, **21**. Assimilatory or dissimilatory nitrate reduction (ANRA or DNRA): 17 or 18 + 19 or 20, **22 & 23**. Nitrite reduction (Denitrification): NO_2_^−^ to NO, **24**. Nitric oxide reduction: NO to N_2_O, **25**. Nitrous oxide reduction: N_2_O to N2. (**e**) Hydrogen metabolism via hydrogenases: **26**. FeFe hydrogenases group A (fermenting and bifurcating), **27**. FeFe hydrogenases group C (H_2_ sensors), **28**. NiFe hydrogenases group 1 (H_2_ oxidation), **29**. NiFe hydrogenases group 2 (H_2_ oxidation), **30**. NiFe hydrogenases group 3b (bidirectional), **31**. NiFe hydrogenases group 3c (bidirectional), **32**. NiFe hydrogenases group 3d (bidirectional). Paired colored bars above any two given boxplots with the same color and at the same level indicate statistically significant differences between those two floodplains (two-way ANOVA).

We also considered the distribution of functions that were detected in < 25% of genomes (**Fig. 4b-e**). Of the remaining C1 transformations examined, methanol oxidation to formaldehyde was found in a median of ~10% of the genomes. Of the sulfur transformations, sulfite (SO_3_^−2^) oxidation to sulfate (SO_4_^−2^), and thiosulfate (S_2_O_3_^−2^) oxidation without sulfur (S^0^) deposition and thiosulfate oxidation with sulfur deposition were most prevalent. For reactions involving hydrogen consumption or formation, genes encoding group 1 NiFe hydrogenases (likely used for H_2_ oxidation) were found in a higher proportion of genomes than any other types of hydrogenases.

Nitrogen transformations were studied individually and as part of the nitrogen cycle. We found the capacity for nitrate (NO_3_^−^) use as a terminal electron acceptor in dissimilatory NO_3_^−^ reduction in a substantially lower proportion of genomes (2 - 10%) than the capacity to use O_2_ as a terminal electron acceptor. Of the reactions involved in nitrification, namely ammonia oxidation, hydroxylamine oxidation and nitrite (NO_2_^−^) oxidation, genomes encoding the oxidation of nitrite via nitrite oxidoreductase (NXR) were more common than genomes encoding the first two steps. The capacity for NO_3_^−^ reduction as part of denitrification (NapAB or NarGHK) was encoded by far fewer genomes than NO_2_^−^ reduction (which can be carried by via multiple enzymes, including NirK, NirS, NrfAH for dissimilatory nitrite reduction or NirBD for assimilation). Fewer genomes are predicted to encode the capacity to reduce nitric oxide (NO, the product of nitrite reduction) to nitrous oxide (via NorBC) than genomes with the capacity for nitrous oxide (N_2_O) reduction to N_2_. Overall, the most prevalent genomically encoded function was nitrite reduction, and capacities for consecutive nitrogen cycling steps were typically encoded in multiple different genomes. In other words, there is evidence to support the prevalence of metabolic handoffs ^7^ in the nitrogen cycle.

We identified functions that were significantly enriched (FDR ≤ 0.05; hypergeometric test) in the core floodplain microbiome (a subset of ISA group 7) and found that the capacities to use O_2_ as a terminal electron acceptor, to perform aerobic CO or other small molecule oxidation, and thiosulfate oxidation (both with and without sulfur deposition) were enriched in these organisms.

### Environmental factors as drivers of function distribution across and within floodplains

Environmental variables (Supplementary Figure 4) may explain in part the patterns of enrichment of genomically encoded functions described above. We first looked into correlations involving the following variables: total carbon (TC), total organic carbon (OC), total inorganic carbon (IC), total nitrogen (TN), organic carbon to nitrogen ratio (OC:N), distance of a sample to the river (Dist. to river), easting and northing (cartesian coordinates for position on the floodplain), distance to the inner bank edge (from here on: toe distance) and distance to middle of the meander-bound floodplain as alternative measures of position on the floodplain (Supplementary Figure 5), topographic position index (TPI; as a proxy for the likelihood a site would be flooded during periods of high discharge or snowmelt), and elevation. Statistical analysis indicated that TC, OC, TN, and OC:N were all highly correlated with each other across the same set of metagenomic samples (Supplementary Figure 6) and their individual effects were not possible to disentangle. Thus, we chose either TC or OC for downstream analyses. Given the Northwest to Southeast orientation of the watershed, elevation, Easting, and Northing were all highly correlated with floodplain (G vs L vs Z), so only floodplain was included as a categorical variable. In summary, TC, floodplain, IC, TPI, distance to the river and toe distance were the variables evaluated with the fourth corner method ^8^ to assess the response of each function at the gene level to the selected environmental or soil chemistry and GIS variables (see **Methods**).

A group of biogeochemical transformations (gene level) displayed some correlation with environmental variables, particularly with individual floodplains (**Fig. 5a**). Genome abundances were used as proxies for abundance of functions each genome encoded. The upstream floodplain G was positively correlated with thiosulfate oxidation (with and without S deposition), sulfite oxidation, sulfide oxidation, O_2_ as a terminal electron acceptor, aerobic CO or other small molecule oxidation, and acetate metabolism. Only N_2_O reduction was positively correlated with the middle floodplain L. The downstream floodplain Z was positively correlated with H_2_ oxidation via group 1 NiFe hydrogenases (a function that was negatively correlated with upstream floodplain G). Most sulfur compound transformations, as well as nitrite reduction, aerobic respiration and acetate metabolism were negatively correlated with floodplain Z (**Fig. 5a**).

**Figure 5.**
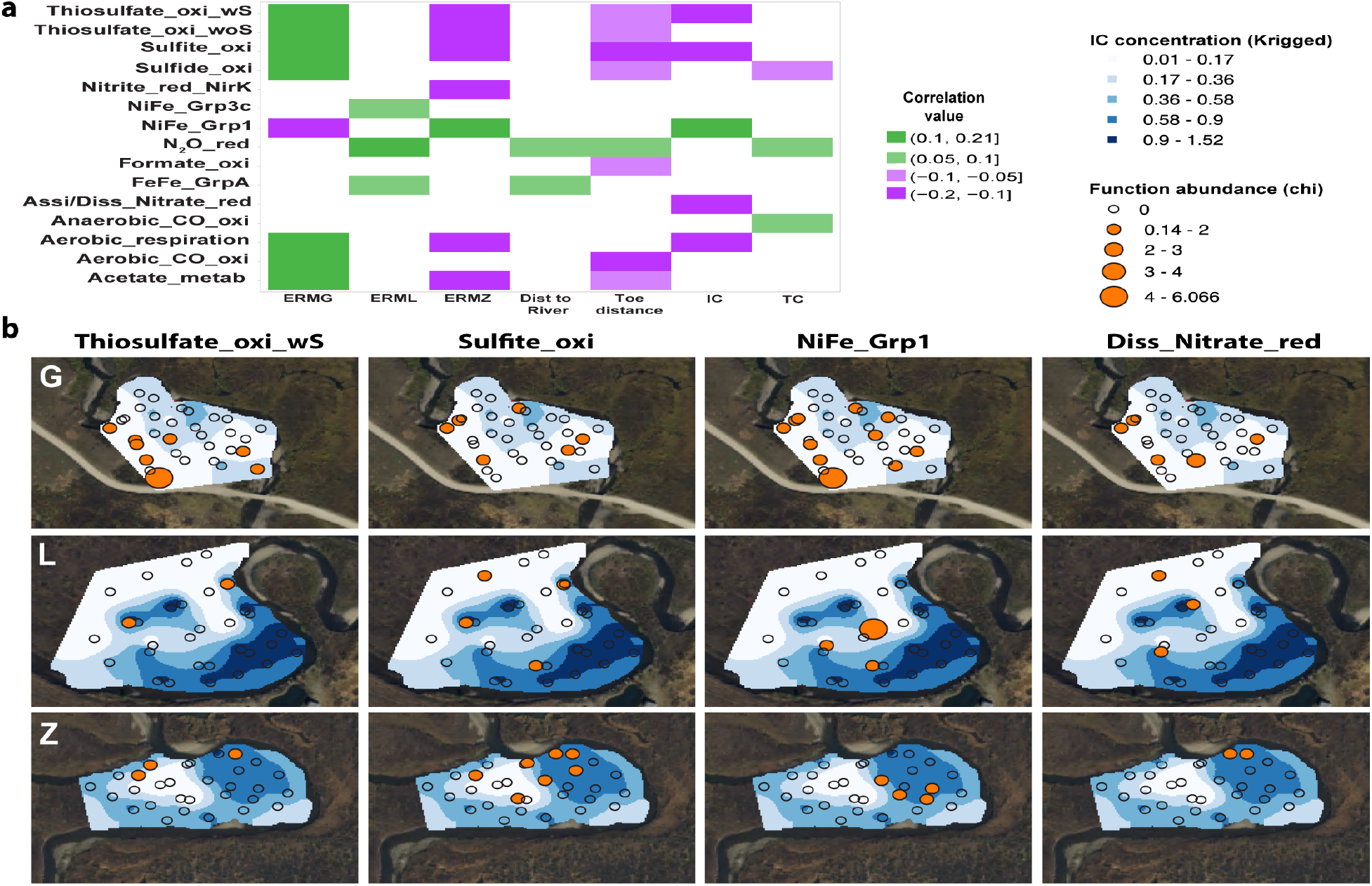
Function abundance and its correlation with environmental variables. **(a)** Significant positive (green) or negative (violet) correlations between environmental variables (bottom) and biogeochemical transformations (left) identified by a fourth corner analysis. **(b)** Abundance of genomes encoding functions positively correlated with inorganic carbon concentrations (IC; %).

Overall, genomes with the capacity for aerobic respiration and sulfur compound oxidation are most prevalent towards the headwaters (floodplain G), and within this floodplain sulfur compound oxidation apparently is associated with low IC (**Fig. 5b**). Within the downstream meander where aerobic respiration is least prominent in the genomes, bacteria able to oxidize H_2_ via Group 1 NiFe hydrogenases appear correlated with somewhat elevated concentrations of IC (**Fig. 5b**).

### Potentially active genes encoding key biogeochemical transformations in the riparian zone

To determine whether key functions encoded in the genomes were transcriptionally active at the time of sampling (early September 2016; during base flow conditions like previous year), we re-sampled floodplain L for metatranscriptomics and metagenomics. This floodplain was chosen among the three because it shared the majority of organisms detected in 2015 with the other two floodplains.

Considering potential differences between the two years, metatranscriptomic reads were mapped to a dereplicated genome set at the species level (95% ANI), which comprised 215 genomes reconstructed from samples collected in 2015 and 2016. We calculated transcript counts using read pairs mapped to predicted open reading frames (ORFs) with at least 95% nucleotide identity (see **Methods**). The highest median transcript counts were observed for Nitrospirae and Betaproteobacteria, followed by Candidatus Latescibacteria and Eisenbacteria-like bacteria, Rokubacteria, and Deltaproteobacteria (**Fig. 6a**). We also evaluated the number of reads mapping to genes encoding key functions and determined what percentile in the distribution of transcription levels each gene fell in.

**Figure 6.**
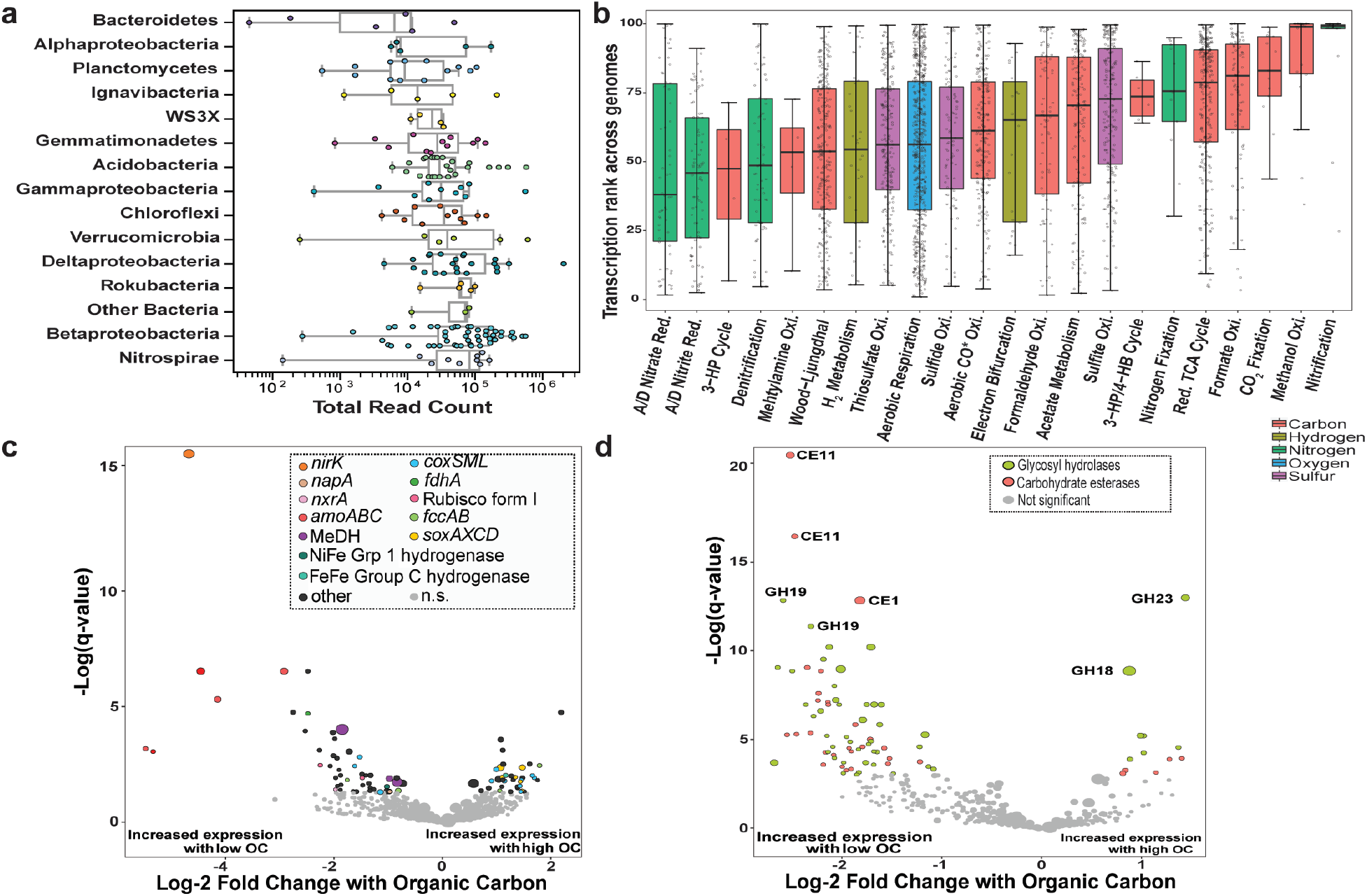
Analyses of transcription from samples collected in 2016 mapped to the species level representative genome set. **(a)** Transcription activity of genomes grouped by phylum or class based on total transcript read counts mapped to the representative genomes. Other Bacteria: Candidatus Latescibacteria and Eisenbacteria-like bacteria. Phylogeny of the genomes at the species level was confirmed based on a concatenated ribosomal proteins tree (Supplementary Data 2). **(b)** Average transcription percentile for all genes encoding enzymes involved in a given biogeochemical transformation in each representative genome across all 2016 metatranscriptomes. (**c**) Differentially transcribed genes in response to soil OC. Statistically significant (DESeq2; q < 0.05) genes are colored by function and not significant (n.s.) genes are in grey. (**d**) Differentially transcribed genes encoding CAZY in response to soil OC.

Key genes involved in potentially active biogeochemical transformations with a median transcription > 75^th^ percentile of all the genes transcribed in a given genome included *amoCAB*, and *nxrAB*, involved in nitrification. The *amoCAB* genes for aerobic ammonia monooxygenase were found in the 90^th^ percentile of transcribed genes across genomes. However, these genes were present in very few genomes (one Nitrospirae and two Thaumarcheota). Similarly present in few genomes and also highly transcribed were genes involved in CO_2_ fixation, specifically RuBisCO forms I and II and enzymes in the reductive TCA cycle (OFOR and citrate lyase). Other highly transcribed genes were for methanol oxidation to formaldehyde (*xoxF, mxaF*) and formate oxidation (*fdhAB, fdoG, fdhF, fdwA, fdsD, fdwB*) as part of C1 metabolism.

Functions enriched in the core floodplain microbiome (aerobic CO or other small molecule oxidation, thiosulfate oxidation, and the ability to use O_2_ as a terminal electron acceptor via *coxABCD, cydAB or ccoN*) and functions that displayed some degree of correlation with environmental variables in gene abundance (*e.g.*, sulfite oxidation via *sat* and *aprAB* or *dsrAB* and sulfide oxidation via *fccAB*) were most often between 50^th^-75^th^ percentile of transcribed genes per genome. Surprisingly given their prominence in genomes, genes involved in nitrogen cycling such as *narGHI or napA* and especially *nrfAH* responsible for dissimilatory nitrite reduction, *nirK* and *nosZ* responsible for some denitrification steps, displayed transcription levels only between the 35^th^ - 50^th^ percentiles (**Fig. 6b**).

We tested for differential transcription levels in response to changes in environmental variables (Supplementary Table 7) using DESeq2 ^9^. Of the four environmental variables that were highly correlated with each other (TC, OC, TN, OC:N; either positively or negatively Supplementary Figure 7), we observed the strongest differential gene expression in response to OC (**Fig. 6c**). In samples with higher concentrations of OC, genes involved in the Sox pathway for thiosulfate oxidation (*soxAX* and *soxCD*) and those involved in aerobic CO or other small molecule oxidation (*coxLMS*) were highly transcribed. More specifically, transcripts mapped to one *coxL* form I gene (true carbon monoxide dehydrogenase, CODH) and the rest mapped to four other *coxL* form II genes (carbon monoxide-like dehydrogenase). The form I transcripts were correlated with high OC, and the form II transcripts with both low (1 hit) and high OC (3 hits).

In samples with low OC, highly transcribed genes included those involved in methanol oxidation (*xoxF, mxaF*), formate oxidation (*fdhA*), sulfide oxidation (*fccAB*), hydrogen metabolism (NiFe Grp1), ammonia oxidation (*amoABC*), nitrite oxidoreduction (*nxrA*), nitrate reduction (*napA*), and nitrite reduction (*nirK*). Samples with low OC also have low TN, which may in part be attributed to high activity of microbial nitrification followed by denitrification, with denitrification reliant on consumption of OC.

As might be expected, RuBisCO form I was highly transcribed under conditions of low OC. Activity of the Calvin Benson Bassham (CBB) pathway for CO_2_ fixation is linked to Betaproteobacteria, Deltaproteobacteria, Gammaproteobacteria, and NC10, some of which have metabolisms fueled by oxidation of intermediate sulfur compounds (*e.g.,* sulfide or thiosulfate oxidation). The observations reveal a potentially important source of organic carbon in some floodplain soils. In terms of overall transcriptional activity, autotrophic pathways may not be expressed but the organisms may otherwise be highly transcriptionally active. In fact, mostly organisms from the phyla Thaumarchaeota, Rokubacteria, NC10 and Nitrospirae were active under low OC conditions. Betaproteobacteria and Acidobacteria were transcriptionally active under conditions of higher OC.

We also investigated the potential for organic matter degradation through transcription of genes encoding carbohydrate-active (CAZY) enzymes (Supplementary Table 8) ^10^. We narrowed our search to CAZY enzyme types that were present in at least 60% of the genomes, as a proxy for widespread distribution in the floodplain soil microbial community. The most abundantly transcribed CAZY genes were in the glycosyl hydrolase (GH) and carbohydrate esterase (CE) classes. In general, the highly transcribed enzymes in the CE class use hemicellulose and amino sugars as substrates, resulting in acetate as a byproduct. Acetate could be utilized by many floodplain organisms, considering the prevalence of genes involved in acetate metabolism among the genomes. Enzymes in the GH class use cellulose, pectin, chitin and starch as substrates, releasing a variety of sugars as byproducts, which can be utilized for central metabolism during growth. We then tested for CAZY differential expression in response to changing concentrations of organic carbon. 12 CAZYs (both GH and CEs) expressed by three strains of Betaproteobacteria increased in expression in samples with high OC. Many of the same classes of GH and CE displayed high levels of transcription correlated with both low and high OC levels (*e.g.*, GH23, GH28, CE4, and CE11 **Fig. 6d**), although gene expression by three strains of Betaproteobacteria correlated with high levels of OC. Similar CAZY enzymes were expressed by NC10, Nitrospirae, and Rokubacteria, the same organisms commonly associated with high levels of transcription under low OC.

## Discussion

Biogeochemical processes modulate C, S, and N exports from watersheds, including the East River ^11^. Important questions relate to the sources and sinks of these compounds and the biological controls on them. Some data indicate that a subset of the organic carbon in sediments from East River floodplains derives from the shale, although plants are the obvious central source for fixed carbon in areas of more developed soils ^12^. CO_2_ fixation genes were relatively rarely detected in the bacterial genomes, which might be interpreted to support this deduction. However, genes for CO_2_ fixation in a few organisms were very highly transcribed, indicating at least periodic inputs of microbially-produced organic carbon into riparian zone soils. Spatially, high activity of genes involved in CO_2_ fixation was correlated with low organic carbon concentrations in soil. Many organisms predicted to rely on CO_2_ fixation as their main carbon source are aerobic chemolithoautotrophs that oxidize inorganic compounds (e.g., NH_3_^+^, NO_2_^−^, S^0^, H_2_S, H_2_, CO, S_2_O_3_) as a source of energy. Thus, we infer significant linkages amongst these key element nutrient cycles.

Low concentration of organic carbon also correlated with high activity of genes involved in methanol oxidation. Methanol results from the breakdown of plant material, such as pectin and lignin, and the activity of methanol dehydrogenases may be indicative of decomposed organic matter. Similarly, low concentration of organic carbon correlated with high activity of genes involved in sulfide and H_2_ oxidation, nitrification and interconversion of nitrite and nitric oxide. The organisms responsible for these reactions are primarily autotrophs.

Interestingly, the capacities for thiosulfate oxidation/elemental sulfur formation, sulfite oxidation and H_2_ oxidation, as well as assimilatory or dissimilatory nitrate reduction to ammonia (ANRA or DNRA) were patchily spatially distributed (**Fig. 4b**), possibly localized by lower inorganic carbon concentrations. Further, genes for sulfur compound oxidation were more prominent in genomes of organisms from the upstream floodplain, which is closer to the adjoining hill, possibly reflecting higher inputs of intermediate sulfur compounds from rock weathering reactions in the headwater compared to downstream regions. The sources of thiosulfate could be weathering of detrital grains of shale-associated pyrite and/or reoxidation of microbially-produced sulfide in anoxic OC-rich regions of the soil or underlying river sediment. Closer proximity to igneous intrusives in the upstream part of the drainage (*e.g.,* near Floodplain G) leads to greater incidence of pyrite-bearing shales. It has been shown previously that hydrological connectivity can shape microbial activity, with low connectivity linked to higher abundance of genes involved in sulfur metabolism ^13^. In the East River, sulfur compounds may be redistributed from upstream to downstream regions, but the degree of hydrologic connectivity within and across floodplains is uncertain and varies dramatically over the course of the year ^14^. Additionally, these shallow soils may only be hydrologically connected to the river during high water flood events or through vertical transport.

By contrast, high OC levels correlated with high activity of genes involved in oxidation of CO (form I CODH), and other small carbon compounds (form II or other subtypes), which may be substrates for carbon monoxide dehydrogenases ^5^. CO may be sourced from the atmosphere, by thermochemical, photochemical, and chemical degradation of organic matter in soils and marine sediments, and from biological production by microbes, leaves, roots and animals ^15^. East River CO oxidizers are most likely carboxydovores that require organic carbon to grow, even though they can oxidize CO at atmospheric levels (*i.e.,* they use a high affinity form I CODH) ^16, 17^. This is in contrast to carboxydotrophs that grow with CO as the sole energy and carbon source and require CO at greater than atmospheric concentrations (for a low affinity form I carbon monoxide dehydrogenase (CODH); ^15^). Additionally, form II CO dehydrogenases seem to play a key role in this ecosystem, although very little is known about their actual function. Detection of a high prevalence of CODH and CODH-like enzymes echoes results from a grassland soil system Diamond, et al. ^5^, reinforcing the suggestion that small carbon compounds such as plant exudates, may be an important carbon currency under some conditions.

High organic carbon levels also correlated with high activity of genes involved in thiosulfate oxidation, and carbohydrate esterases and glycosyl hydrolases such as GH23 (lysozyme) and GH18 (chitinase). These GHs would be required for organic matter degradation at locations of higher carbon availability, where presumably plants and fungi are more abundant. Bacteria that degrade plant biomass are also known to employ catabolite repression of CAZy enzymes ^18 35,17^, perhaps explaining the lower diversity of CAZys under these conditions. Different variants of these carbohydrate-active genes were highly expressed in a variety of taxa including Rokubacteria, Nitrospirae and NC10 in soil with low organic carbon, where diverse carbon sources must be exploited for survival.

Many watershed ecosystems are limited by access to biologically available nitrogen, the important sources of which are likely to be shale bedrock weathering ^19^, atmospheric deposition ^20^, and nitrogen fixation. A complex interplay of biological processes impact nitrogen speciation and bioavailability, including ammonia oxidation (nitrification), denitrification to N_2_, and nitrite assimilation via ANRA or DNRA. The nitrogen budget can be addressed by direct measurement of inputs, plant-associated inventories, and the concentration of inorganic and organic nitrogen compounds exported from the watershed via rivers ^21, 22^. By comparing these numbers, it may be possible to estimate the fraction of the bioavailable nitrogen that is lost from the system via loss as N_2_ and trace gases. What is missing from this analysis is an estimate of the degree to which nitrogen compounds are recycled, the role of riparian zone soils in these processes, and the potential for subsurface storage of nitrogen compounds in microbial biomass.

Using genome-resolved metagenomics we identified the capacity for nitrogen fixation and ammonia oxidation to nitrite and nitrate in relatively few organisms, yet the metatranscriptomic data show these to be highly active functions. Thus, we infer important microbial contributions to reservoirs of oxidized nitrogen compounds in riparian zone soils, with the potential to substantially augment inputs from atmospheric deposition and bedrock weathering. Genes involved in nitrite reduction (dissimilatory nitrate reduction to ammonium or denitrification to N_2_), while abundant in comparison to other capacities for nitrogen transformation, displayed surprisingly low levels of transcription at the time of sampling. Over the course of the year, the fluctuating water table and periodic flooding should provide environmental niches for both obligately aerobic, and anaerobic processes. Furthermore, the soil oxidation state and the carbon to nitrate ratio, particularly in nitrogen-limited systems, may favor DNRA over denitrification ^23^. The current study was conducted during base flow conditions (both years), well after the snowmelt season, when high river discharge induces flooding and therefore anoxic conditions. In the meander-bound floodplains, snowmelt-derived flow in this ecosystem persists well into the year ^12^, so shallow soils may be flooded long after discharge levels drop. The results raise the possibility of coupling of nitrification and dissimilatory nitrate pathways on a temporal basis, under baseflow conditions (when nitrification is dominant), or under snowmelt conditions (when dissimilatory processes occur). High net nitrification has been reported for riparian zones when the water table is below −30 cm ^24^, accordingly the water table in floodplain L was observed to be below this level in September 2016. Overall, re-assimilation of nitrogen as ammonium may be important in this ecosystem, particularly if nitrogen limited.

An important result from the current study was that there appears to be a core floodplain microbiome composed of specific bacterial species from Betaproteobacteria, Gammaproteobacteria, Deltaproteobacteria, Nitrospirae, Candidatus Latescibacteria, and Rokubacteria; and all of these groups were transcriptionally active at the time of sampling. Many of the clusters of related genomes are relatively distantly related to previously described bacterial types. Thus, we conclude that many of the most abundant taxa in these riparian zone soils are organisms that have, until now, remained essentially outside of the range of scientific investigations. Importantly, capacities for aerobic respiration, aerobic oxidation of CO and other small molecules, as well as thiosulfate oxidation with formation of elemental sulfur, were enriched in the core floodplain microbiome. Notably, the most abundant functions of the core microbiome were only moderately transcribed at the time of sampling.

In general, we found that gene and organism abundances do not predict transcription levels. The *in situ* transcription data revealed the potentially very high importance of rare genes and organisms. However, it is important to note that transcript datasets are a snapshot from a moment in time, and that transcription patterns will vary across seasons and maybe even daily. Notably, our analyses showed organismal and functional overlap in microbial communities found both within and across the three floodplains over two consecutive years (~15% of species in common, despite the very high diversity and complexity of the soils). Thus, in contrast to potentially substantial transcriptome variability, gene inventories reflect metabolic potential that likely remains fairly constant throughout the year. Thus, we conclude that, at the watershed scale, meander-bound regions of floodplain soils are “functional zones” that likely predict biogeochemical transformations along the riparian corridor, thereby providing broadly generalizable inputs to ecosystem models.

## Methods

### Study site and samples collection

The East River (ER) watershed has been described elsewhere ^3^. In brief, the ER watershed is a 300 km^2^ area largely underlain by marine shales of the Cretaceous Mancos formation located in the Elk Mountains in west-central Colorado. The ER is a headwaters catchment in the Upper Colorado River basin, with an average elevation of 3350 m. At about 62 km long, the ER traverses an elevational gradient that includes alpine, subalpine, and montane life zones as a function of stream reach. The average annual temperature is ~ 0 °C, with long cold winters and short cool summers, and the majority of precipitation is received in the form of snow ^25^.

The sampling sites are located across an altitudinal gradient followed by the river (~2700 – 2900 m). The floodplain at the highest elevation is located ca. 6 km from the headwaters, nearby Gothic, Colorado, site of the Rocky Mountain Biological Laboratory (RMBL, **Fig. 1**). Therefore, samples collected from this site were named East River Meander-bound floodplain G (ERMG). The second site was located ca. 8 km downstream of Gothic, among a series of floodplains, one of which is situated adjacent to an intensive research site of the Watershed Function SFA ^3^. This floodplain stands out because of its larger size, and samples were named ERML (L for large). The third site was located ca. 18 km downstream of Gothic and just upstream of the confluence with Brush Creek. Samples from this site were named ERMZ, with the stream reach between ERML and ERMZ being characterized by a relatively low gradient with high sinuosity.

In September 2015, during base flow conditions, two series of perpendicular transects were laid out at each site. Each set of transects comprised four transects that were parallel between them (**Fig. 1**). One set of transects were approximately North to South (T1-T4) and the other set of transects were East to West (T5-T8). The starting point of each transect was designated “0 m” and the location of the other sites along the transect was relative to the point of origin. A Trimble Geo 7X GPS was used to determine the exact location of each site along the transects with an accuracy of 0.5 m. The distance (in meters) of each sample to the point of origin was included in the sample name, which comprised the initials of the study area (ER), the initials for each meander-bound floodplain (*i.e.,* MG, ML or MZ), the transect number (*i.e.,* T1-T8), and the distance in meters from the first sample collected at the start point (*e.g.,* 19 m). We sampled an area ~ 4,600 m^2^ in floodplain G, ~ 8,000 m^2^ in floodplain L, and ~ 5,400 m^2^ in floodplain Z.

Four soil samples from the 10-25 cm (± 1-2 cm) soil depth interval were collected in the span of 10 days along each one of the eight transects, for a total of 32 samples per floodplain. Each site was cleared of grasses and other vegetation with clippers, and the first ~ 10 cm of soil was removed with a sterile shovel. Soil samples were collected using sterile tools, including a soil core sampler and 7.6 × 15.2 cm plastic corer liners (AMS, inc), stainless-steel spatulas, and Whirl-pak bags. Samples were immediately stored in coolers for transportation to RMBL, where samples were prepared for archival and transportation to the University of California, Berkeley. Soil cores were broken apart and manually homogenized inside the Whirl-pak bags. Subsamples for chemical analyses, DNA extractions, and long-term archival were obtained inside a biosafety cabinet, kept at – 80 °C, transported in dry ice, and stored at – 80 °C at the University of California, Berkeley.

In September 2016, another round of samples collection was conducted at floodplain L for metagenomics, metatranscriptomics, and chemical analyses. A subset of 19 out of the 32 sampling sites from the previous year was targeted, and a subset (15) of those was also selected for metatranscriptomics (Supplementary Table 1). Given that floodplain L was the site with the lowest total number of draft genomes recovered in 2015, we added new sites closer to the original sites with the intent of increasing this number by leveraging differential coverage across samples ^26^. Four new sites located **i**n **b**etween the original **t**ransects (denominated ERML**IBT**) and two sites adjacent to ERMLT660 (ERMLT660_1 and ERMLT660_2) were sampled. Additionally, samples were collected from above the water table (approximately below 40-50 cm from the surface) at a depth of 32-47 cm (± 4-6 cm) from three sites (ERMLT200, ERML231 and ERML293) along T2. Samples from the 11-25 cm (± 1-1 cm) soil layer were obtained following the same protocol as the previous year, with the exception that subsamples for RNA sequencing were preserved *in situ*. Once the soil cores were transferred to a Whirl-pak bag, they were manually homogenized inside the bags. Eight grams (8 g) of soil were collected using sterile stainless-steel spatulas directly into 50 mL sterile falcon tubes containing 20 mL of LifeGuard Soil Preservation Solution (formerly MoBio) for RNA preservation. The samples were mixed by hand to saturation with the LifeGuard solution, stored in a chilled cooler for transportation to RMBL and later stored at −80 °C.

### Soil chemistry

Total carbon (TC) and total inorganic carbon (TIC) were analyzed using a Shimadzu TOC-VCPH analyzer equipped with a solid sample module SSM-5000A (Shimadzu Corporation, Japan). Total organic carbon (TOC) was obtained from the difference between TC and TIC. For TC quantification, a subsample of the dried solids was weighed into a ceramic boat and combusted in a TC furnace at 900 °C with a stream of oxygen. To ensure complete conversion to CO_2_, the generated gases are passed over a mixed catalyst (cobalt/platinum) for catalytic post-combustion. The CO_2_ produced is subsequently transferred to the NDIR detector in the main instrument unit (TOC-VCSH). Quantification of the inorganic carbon was carried out in a separate IC furnace of the module. Phosphoric acid is added to the sample and the resulting CO_2_ is purged at 200 °C and measured.

Total nitrogen (TDN) was analyzed using a Shimadzu Total Nitrogen Module (TNM-1) coupled to the solid sample module (SSM-5000A) and TOC-VCSH analyzer (Shimadzu Corporation, Japan). TNM-1 is a non-specific measurement of TN. All nitrogen species in samples were combusted at 900 °C, converted to nitrogen monoxide and nitrogen dioxide, then reacted with ozone to form an excited state of nitrogen dioxide. Upon returning to the ground state, light energy is emitted. Then, TN is measured using a chemiluminescence detector.

### DNA extraction and sequencing

Genomic DNA was extracted from ~10 g of thawed soil using Powermax Soil DNA extraction kit (Qiagen) with some minor modifications as follows. Initial cell lysis by vortexing vigorously was substituted by placing the tubes in a water bath at 65 °C for 30 minutes and mixing by inversion every 10 minutes to decrease shearing of the genomic DNA. After adding the high concentration salt solution that allows binding of DNA to the silica membrane column used for removal of chemical contaminants, vacuum was used instead of multiple centrifugation steps. Finally, DNA was eluted from the membrane using 10 mL of the elution buffer (10 mM Tris buffer) instead of 5 mL to ensure full release of the DNA. DNA was precipitated out of solution using 10 mL of a 3 M sodium acetate (pH 5.2) and glycogen (20 mg/mL) solution and 20 mL 100% sterile-filtered ethanol. The mix was incubated overnight at 4 °C, centrifuged at 15,000 x *g* for 30 minutes at room temperature, and the resulting pellet was washed with chilled 10 mL sterile-filtered 70% ethanol, centrifuged at 15,000 x *g* for 30 min, allowed to air dry in a biosafety cabinet for 15-20 minutes, and resuspended in 100 μL of the original elution buffer. Genomic DNA yields were between 0.1 – 1.0 μg/μL except for two samples with 0.06 μg/μL. Power Clean Pro DNA clean up kit (Qiagen) was used to purify 10 μg of DNA following manufacturer’s instructions except for any vortexing was substituted by flickering of the tubes to preserve the integrity of the high molecular weight DNA. DNA was resuspended in the elution buffer (10 mM Tris buffer, pH 8) at a final concentration of 10 ng/μL and a total of 0.5 μg of genomic DNA. DNA was quantified using a Qubit double-stranded broad range DNA Assay or the high-sensitivity assay (ThermoFisher Scientific) if necessary. Additionally, the integrity of the genomic DNA was confirmed on agarose gels and the cleanness of the extracts tested by absence of inhibition during PCR. For samples collected the following year, DNA was co-extracted with RNA (see next section), in addition to extracting subsamples (10 g of soil) from the same core following the extraction protocol described above (Supplementary Table 1).

Clean DNA extracts and co-extracts were submitted for sequencing at the Joint Genome Institute (Walnut Creek, CA), where samples were subjected to a quality control check. Two of the 96 samples from 2015 failed QC and thus were not sequenced (ERMZT233 and ERMZT446), and four samples were sequenced ahead of the others (ERMLT700, ERMLT890, ERMZT100, and ERMZT299). Ten out of 15 of the DNA co-extracts from 2016 failed QC due to low DNA yields and were not sequenced either. Sequencing libraries for the first four samples were prepared in microcentrifuge tubes. 100 ng of Genomic DNA was sheared to 600 bp pieces using the Covaris LE220 and size selected with SPRI using AMPureXP beads (Beckman Coulter). The fragments were treated with end-repair, A-tailing, and ligation of Illumina compatible adapters (IDT, Inc) using the KAPA Illumina Library prep kit (KAPA biosystems). Libraries for the rest of the samples were prepared in 96-well plates. Plate-based DNA library preparation for Illumina sequencing was performed on the PerkinElmer Sciclone NGS robotic liquid handling system using Kapa Biosystems library preparation kit. 200 ng of sample DNA was sheared to 600 bp using a Covaris LE220 focused-ultrasonicator. The sheared DNA fragments were size selected by double-SPRI and then the selected fragments were end-repaired, A-tailed, and ligated with Illumina compatible sequencing adaptors from IDT containing a unique molecular index barcode for each sample library.

All the libraries were quantified using KAPA Biosystem’s next-generation sequencing library qPCR kit and a Roche LightCycler 480 real-time PCR instrument. The quantified libraries were then multiplexed with other libraries, and the pool of libraries was prepared for sequencing on Illumina HiSeq sequencing platform utilizing a TruSeq paired-end cluster kit, v4, and Illumina’s cBot instrument to generate a clustered flow cell for sequencing. Sequencing of the flow cell was performed on the Illumina HiSeq 2500 sequencer using HiSeq TruSeq SBS sequencing kits, v4, following a 2×150 indexed run recipe.

### RNA-DNA co-extraction and sequencing

Total RNA was extracted from a subset of 15 samples using the RNA PowerSoil Total RNA isolation kit (Qiagen). Soil samples (8 g) preserved in LifeGuard solution (Qiagen) were thawed on ice and centrifuged at 2,500 x *g* for 5 minutes to collect the soil at the bottom of the tubes. As a supernatant, the LifeGuard solution was extracted from the tubes and aliquoted into three 15 mL conical tubes that were used to transfer three separate 2 g subsamples for later use. The remaining 2 g were split in half into two of the kit’s bead tubes with pre-aliquoted bead solution (to disperse the cells and soil particles). The lysis solution (SR1) and the non-DNA organic and inorganic precipitation solution (SR2) were not added to the bead tube until all the subsamples to be processed in a given day had been aliquoted. Subsamples were kept at −20 °C before transferring them to a −80 °C freezer for permanent storage. The remainder of the extraction was carried out following the manufacturer’s instructions. An RNA PowerSoil DNA elution accessory kit was used to co-extract DNA from the RNA capture columns, which was quantified as previously described. A DNase treatment was performed in all the RNA extracts with a TURBO DNA-free kit (Ambion) using 4 U of TURBO DNase at 37 °C for 30 minutes. The absence of DNA was tested by PCR with universal primers to the SSU rRNA gene, and the integrity of the RNA was checked using a Bioanlayzer RNA 6000 Nano kit following the manufacturer’s instructions. Total RNA was quantified before and after DNase treatments using a Qubit high-sensitivity RNA assay (ThermoFisher Scientific). One of the RNA extracts (ERMLT590) did not yield enough RNA for sequencing.

Total RNA and DNA co-extracts were submitted for sequencing at the Joint Genome Institute in Walnut Creek, CA, where samples were subjected to a quality control check. rRNA was removed from 1 μg of total RNA using Ribo-Zero(TM) rRNA Removal Kit (Illumina). Stranded cDNA libraries were generated using the Illumina Truseq Stranded mRNA Library Prep kit. The rRNA depleted RNA was fragmented and reversed transcribed using random hexamers and SSII (Invitrogen) followed by second strand synthesis. The fragmented cDNA was treated with end-pair, A-tailing, adapter ligation, and 8 cycles of PCR. For low input extracts, rRNA was removed from 100 ng of total RNA using Ribo-Zero(TM) rRNA Removal Kit (Illumina). Stranded cDNA libraries were generated using the Illumina Truseq Stranded mRNA Library Prep kit. The rRNA depleted RNA was fragmented and reversed transcribed using random hexamers and SSII (Invitrogen) followed by second strand synthesis. The fragmented cDNA was treated with end-pair, A-tailing, adapter ligation, and 10 cycles of PCR. The prepared libraries were quantified using KAPA Biosystem’s next-generation sequencing library qPCR kit and run on a Roche LightCycler 480 real-time PCR instrument. The quantified libraries were then multiplexed with other libraries, and the pool of libraries was prepared for sequencing on the Illumina HiSeq sequencing platform utilizing a TruSeq paired-end cluster kit, v4, and Illumina’s cBot instrument to generate a clustered flow cell for sequencing. Sequencing of the flow cell was performed on the Illumina HiSeq 2500 sequencer using HiSeq TruSeq SBS sequencing kits, v4, following a 2 × 150 indexed run recipe.

### Metagenomes assembly and annotation and ribosomal protein L6 analysis

Methods used for 2015 and 2016 metagenomes assembly and annotation are described elsewhere ^27^. In brief, after quality filtering, reads from individual samples were assembled separately using IDBA-UD v1.1.1 with a minimum k-mer size of 40, a maximum k-mer size of 140 and step size of 20. Only contigs > 1Kb were kept for further analyses. Gene prediction was done with Prodigal v2.6.3 in meta mode, annotations obtained using USEARCH against Uniprot, Uniref90 and KEGG, and 16S rRNA and tRNAs predicted as described in Diamond et al. ^5^. Reads were mapped to the assemblies using Bowtie2 ^28^ and default settings to estimate coverage. To estimate the number of genomes potentially present across all 94 metagenomes, we used the ribosomal protein L6 as marker gene and RPxSuite (https://github.com/alexcritschristoph/RPxSuite) as described in Olm et al. ^6^.

### Genome binning, curation, and dereplication

Annotated metagenomes from both years were uploaded onto ggKbase (https://ggkbase.berkeley.edu), where binning tools based on GC content, coverage and winning taxonomy ^29^ were used for genome binning. These bins and additional bins that were obtained with the automated binners ABAWACA1 (https://github.com/CK7/abawaca), ABAWACA2, MetaBAT ^30^, Maxbin2 ^31^ and Concoct ^32^ were pooled, and DAStool was used for selection of the best set of bins from each sample as described by Diamond et al. ^5^. Notably, no bins were recovered from sample ERMZT266 by any method.

Genomic bins were filtered based on completeness ≥ 70% of a set of 51 bacterial single copy genes (BSCG) if affiliated with Bacteria and a set of 38 archaeal single copy genes (ASCG); and a level of contamination ≤ 10% based on the corresponding list of single copy genes ^33^. Additionally, bins that were 59-68% complete with a highest taxonomic level defined as Bacteria in ggKbase, or potential members of the candidate phyla radiation (CPR) were kept for further scrutiny. To obtain a set of genomes for visual curation in ggKbase, genomes were dereplicated at 99% ANI across samples located within a given floodplain using dRep with the --ignoreGenomeQuality flag. ^34^. Any assembly error in the dereplicated set was addressed using ra2.py ^35^, and contigs that fell below the 1 Kb length minimum after this step were removed from the bins. At this point, the level of completeness of CPR genomes was confirmed based on a list of 43 BSCG ^7^. Genomes that did not meet the completeness thresholds post-assembly error correction and that were not affiliated with CPR or novel bacteria were removed from the analysis. Considering that bins changed as a result of this process, genes were re-predicted using Prodigal in single mode, reads were mapped to the bins using Bowtie2, and bins were re-imported onto ggKbase. Visual inspection of taxonomic profile, GC content and to a minor extent coverage, allowed us to further reduce contamination. The final set of 248 curated bins from 2015 was dereplicated at 98% ANI this time across floodplains including the --genomeInfo flag to take into account completeness and contamination in the process of representative bin selection. Within this set, genomes ≥90% complete were deemed near-complete (Supplementary Table 2). Eight relatively low coverage genomes fell just below the completeness requirement due to fragmentation after curation to remove possible local assembly errors; these were retained as they represent important taxonomic diversity.

Similarly, genomes reconstructed from floodplain L samples collected in 2016 that passed the completeness (≥ 70%) and contamination thresholds (≤ 10%) were visually inspected and improved in ggKbase. Assembly errors were corrected with ra2.py ^35^, and contigs that fell below the 1Kb length were removed, as well as genomes that did not pass the thresholds for completeness after assembly error correction. Genes were re-predicted using Prodigal in single mode and the final set of curated genomes were imported onto ggKbase.

To determine whether the same species were present in two different years, we pooled the genome set from 2015 and the curated 2016 set and dereplicated using dRep at 95% ANI including the --genomeInfo flag to take into account completeness and contamination in the process of representative bin selection ^34^. In this set of genomes, 13 were reconstructed from a deeper depth (Supplementary Table 3). However, only 3 genomes were unique and the other 10 clustered with genomes reconstructed from the ~10-25 cm depth, indicating overlap between the species found at the two depths. Therefore, we kept these genomes for further analyses.

### Genome metabolic annotation

We carefully chose a set of ecologically relevant proteins that catalyze geochemical transformations related to aerobic respiration, metabolism of C1 compounds, hydrogen metabolism, nitrogen cycling, and sulfur cycling (Supplementary Table 4). Hidden Markov Models (HMMs) for the majority of these proteins were obtained from KOfam, the customized HMM database of KEGG Orthologs (KOs) ^36^. Custom-made HMMs targeting nitrite oxidoreductase subunits A and B (NxrA and NxrB), periplasmic cytochrome *c* nitrite reductase (NirS, cd1-NIR heme-containing), cytochrome *c*-dependent nitric oxide reductase (NorC; cNOR), hydrazine dehydrogenase (HzoA), hydrazine synthase (HzsA), dissimilatory sulfite reductase D (DsrD), sulfide:quinone reductase (Sqr), sulfur dioxygenase (Sdo), ribulose-bisphosphate carboxylase (RuBisCO) form I and form II, and alcohol dehydrogenases (Pqq-XoxF-MxaF) were obtained from Anantharaman et al. ^7^. NiFe and FeFe hydrogenases were predicted using HMMs from Méheust et al. ^37^ and assigned to functional groups following Matheus Carnevali et al. ^29^ (see Phylogenetic Analyses below for tree construction methods; Supplementary Data 3 and 4 and Supplementary Tables 9 and 10). No real group 4 membrane bound NiFe hydrogenases were identified among the East River representative genomes (data not shown). HMMER3 ^38^ was used to annotate the dereplicated sets of genomes following predefined score cutoffs ^36^. A subset (10%) of the hits to all of these HMMs were visually checked to determine whether the cutoffs were appropriate for this dataset as described in Lavy et al. ^39^ and Jaffe et al. ^40^. Only in the case of formate dehydrogenase (FdhA (K05299 and K22516), FdoG/FdhF/FdwA (K00123)) the cutoff was lowered to include additional hits.

For a protein to be considered potentially encoded in the genome, the catalytic subunit and the majority of the accessory subunits had to be detected by the corresponding HMMs at the established cutoffs. The implication for these function definitions is that in some cases even if some subunits that make up an enzyme were detected, the enzyme could have been deemed absent because a key part was missing (Supplementary Table 4). Similarly, pathways that require the activity of multiple enzymes were only detectable if all of the enzymes were present. Only in cases like the Wood-Jungdahl pathway we required the majority of the genes to be present, taking into consideration genome completeness. Furthermore, if multiple enzymes could catalyze a given reaction (*e.g.,* use O_2_ as a terminal electron acceptor) the presence of genes encoding one such enzyme in a genome would be indicative that this capacity was present in the genome. Additionally, if different pathways lead to the same biogeochemical transformation (*e.g.,* CO_2_-fixation), the presence of genes encoding one of those pathways (or key enzymes) was considered as sufficient to indicate its presence (Supplementary Table 4). In a limited number of cases a given pathway may also involve enzymes that are part of central metabolism or that are part of multiple pathways, and in these cases we chose to define presence based on the key catalyst instead of the whole pathway (*e.g.,* RuBisCO in the Calvin Benson pathway).

Carbohydrate active enzymes were predicted using the Carbohydrate-Active enZYmes Database (CAZY; http://www.cazy.org/) ^10^ (version 1.0) (e-value cut-off 1e-20).

### Genome coverage and detection

Reads were mapped to the dereplicated set of bins using Bowtie2 ^28^ and a mismatch threshold of 2% dissimilarity. Calculate_coverage.py (https://github.com/christophertbrown/bioscripts/tree/master/ctbBio) was used to estimate the average number of reads mapping to each genome and the proportion of the genome that was covered by reads (breadth). Genomes with a coverage of at least 0.01 X were considered to be detected in a given sample. The Hellinger transformation was used to account for differences in sequencing depth among samples and determine final genome abundance. To illustrate genome detection across samples we used the ggplot2 package ^41^. Genomes were clustered by average linkage using the Hellinger transformed abundance across samples (from read mapping), and the samples were clustered by Euclidean distance in R ^42^.

### Phylogenetic analyses

Two phylogenetic trees were constructed with a set of 14 ribosomal proteins (L2, L3, L4, L5, L6, L14, L15, L18, L22, L24, S3, S8, S17, and S19). One tree included Betaproteobacteria genomes from this study at the subspecies level (98% ANI) and ~ 1540 reference Betaproteobacteria genomes from the NCBI (Supplementary Figure 2 and Supplementary Data 1). The other tree included the set of 215 genomes dereplicated at 95% ANI and ~ 2,228 reference genomes from the NCBI genome database (Supplementary Data 2). For each genome, the ribosomal proteins were collected along the scaffold with the highest number of ribosomal proteins. A maximum-likelihood tree was calculated based on the concatenation of the ribosomal proteins as follows: Homologous protein sequences were aligned using MAFFT (version 7.390) (--auto option) ^43^, and alignments refined to remove gapped regions using Trimal (version 1.4.22) (--gappyout option) ^44^. Tree reconstruction was performed using IQ-TREE (version 1.6.12) (as implemented on the CIPRES web server ^45^, using ModelFinder ^46^ to select the best model of evolution (LG+I+G4), and with 1000 ultrafast bootstrap ^47^. Taxonomic affiliations were determined based on the closest reference sequences relative to the query sequences on the tree and extended to other members of the ANI cluster. In many cases, the phylogeny was not clear upon first inspection of the tree and additional reference genomes were added if publicly available.

Phylogenetic trees for proteins of interest were reconstructed using the same methods described above, except with different sets of reference sequences. East River homologs in the dimethyl sulfoxide reductase (DMSOR) superfamily such as the catalytic subunit of formate dehydrogenase (FdhA), nitrite oxidoreductase (NxrA), membrane-bound nitrate reductase (NarG; H^+^-translocating), and periplasmic nitrate reductase subunit A (NapA) were confirmed by phylogeny on a tree with reference sequences from Méheust et al. ^37^ (Supplementary Table 11 and Supplementary Data 5). To distinguish form I and form II CODHs and other other subtypes among homologs to K03520 we used Diamond’s et al. ^5^ dataset, which comprises reference sequences from Quiza et al. ^16^ (Supplementary Table 12 and Supplementary Data 6). Similarly, homologs identified using the Pqq-XoxF-MxaF HMM for alcohol dehydrogenases were placed on a phylogenetic tree with reference sequences from Diamond’s et al. ^5^ dataset, comprising references from Keltjens et al. ^48^ and Taubert et al. ^49^. In this tree, all East River homologs were clustered with methanol dehydrogenases (Supplementary Table 13 and Supplementary Data 7) instead of other types of alcohol dehydrogenases. To distinguish between dissimilatory (bi)sulfite reductase oxidative or reductive bacterial types, DsrA and DsrB homologs from individual genomes were concatenated to each other, aligned, and added to a phylogenetic tree with reference sequences from Muller et al. ^50^ (Supplementary Table 14 and Supplementary Data 8).

### Community diversity and composition

Diversity indices for each sample were calculated from the Hellinger transformed abundance table for the genome set at subspecies level (98% ANI) using the vegan package in R ^51^. Species numbers and Shannon diversity per sample were quantified using the specnumber and vegdist functions of vegan respectively (Supplementary Figure 3). An analysis of variance, implemented in the aov function in R, was used to test for significant differences in mean species number and Shannon diversity in relationship to the floodplain samples originated from. No significant differences in group means were detected considering a p-value < 0.05 as significant.

To investigate community composition at the phylum/class level as determined by phylogenetic analysis, the Hellinger-transformed abundance table for the genome set at the subspecies level (98% ANI) was converted to a presence/absence table. The number of samples where each genome was detected was counted and the number of genomes affiliated to a given taxon was summed by sample and plotted in R with ggplot2 ^41^.

### Identification of a core floodplain microbiome

To identify organisms that were a “core” or “shared” set across all sampled sites, we operationally defined a core set as: (1) organisms that were not statistically associated with any specific floodplain using indicator species analysis, and (2) who were detected (displayed ≥ 0.01X coverage) in at least 89 of the 94 total samples (the 90th percentile for this level of presence across all 248 genomes). Indicator species analysis was performed on the log transformed coverage values that were filtered to include only coverage values ≥ 0.01X using the indicspecies package ^52^ in R version 3.5.2 (R core team 2018) ^42^ with 9999 permutations. All p-values for associations of an organism genome with a floodplain or group of floodplains were then subsequently corrected using False Discovery Rate with FDR ≤ 0.05 being considered a significant association. This resulted in 42 genomes that were not statistically associated with any floodplain by ISA and were also detected in ≥ 89 samples (Supplementary Table 5). For visualization of organism abundance profiles in relationship to their membership in the core floodplain microbiome, ISA clusters, and relative to the coefficient of variation of their coverage, Hellinger normalized coverage data was projected onto a two dimensional space using Uniform Manifold Approximation and Projection (UMAP) implemented in the uwot package in R ^53^ using the following parameters: umap(data = coverage_data, n_neighbors = 15, nn_method = “fnn”, spread = 5, min_dist = 0.01, n_components = 2, metric = “euclidian”, n_epochs = 1000).

### Identification of enriched metabolic functions in core floodplain microbiome

Overrepresentation of metabolic functions within the set of genomes comprising the core floodplain microbiome (n = 42) was assessed using hypergeometric testing. The probability of observing the number of genomes in the core floodplain microbiome carrying each of 33 functions, given the total number of genomes with that function across our full genomic dataset (n = 248), was calculated using the phyper function in R. Probabilities calculated across all metabolic functions were corrected for multiple testing using false discovery rate with the p.adjust function in R and with FDR ≤ 0.05 being considered a significant enrichment of a function in the core microbiome.

### Analysis of correlations among environmental variables

Correlations between numeric soil biogeochemical variables across samples were calculated using spearman rank correlation implemented in the rcorr function of the Hmisc package in R (https://github.com/harrelfe/Hmisc). Correlations between variables were then plotted as a correlogram and ordered using hierarchical clustering with Ward’s method using the corrplot package in R ^54^.

### Fourth corner analysis

A rlq-fourth corner analysis was performed on genome abundances, environmental data, and genome metabolic annotations using the R package *ade4* ^55^. Specifically, the pre-Hellinger transformed genome abundance table was used for a correspondence analysis, the selected environmental variables (see *Soil Chemistry* and *GIS*) were used for a Hill-Smith analysis, and the genome metabolic annotations were used for PCA. A randomization test (as described by ter Braak et al. ^56^ and Dray et al. ^57^ was used to test the global significance of the trait-environment relationships. The fourth-corner statistic was then calculated on the same inputs as the rlq analysis with 50,000 permutations and p-value adjustments using the FDR global methods. The results of the rlq-fourth corner analysis were plotted using the ggplot2 package ^41^.

### Metatranscriptomic analyses

To determine differentially transcribed genes, potential levels of activity by phylum or class, most transcribed CAZY, and most transcribed genes among key geochemical transformations, metatranscriptomic reads were mapped using Bowtie2 ^28^ to a set of high-quality draft genomes dereplicated at 95% (see above). Read pairs were then filtered by a minimum identity of 95% to the reference with MAPQ>=2 and total number of mapped read pairs was counted for each gene. Counts for metabolic genes were analyzed with DESeq2 ^9^ to determine differential expression in response to soil organic carbon and p-values were adjusted to correct for multiple hypothesis testing (FDR<0.05).

### GIS

All GIS operations and cartographic visualizations were performed in QGIS v2.12.1 except where otherwise stated. The base remote sensed imagery used was obtained from USDA NAIP (USDA-FSA Aerial Photography Field Office publication date 20171220; 1m ground pixel resolution). Digital terrain model (DTM) at a ground resolution of 0.5 m/pixel was derived by airborne LiDAR data acquired by Quantum Spatial in collaboration with Eagle Mapping Ltd ^58^ (doi:10.21952/WTR/1412542) in 2015. All maps were projected using EPSG:26913 NAD83/ UTM zone 13N. Meander and adjacent river polygons were manually delineated in QGIS. The distance from a sample point to the manually delineated river polygons was calculated using the NNJoin tool. To calculate the sample distances to meander toe, lines were manually drawn between all samples and the meander toe perpendicular to river flow and distances calculated using NNJoin (Supplementary Figure 5). Similarly, to calculate sample distances to the middle of the meander, a line perpendicular to the meander toe line was drawn across the middle of the meander (Supplementary Figure 5). Sample distances to this line were also calculated using NNJoin and samples on the downstream side of the line were converted to negative values to indicate upstream and downstream sides of the meander. TPI is computed from the DTM as the difference between the elevation of a center point and the average elevation measured in the neighboring area (3 by 3 m) ^59^. To display genome abundances as used in the rlq-fourth corner analysis, filtered abundance values were chi-square transformed in R using the *decostand* in the vegan package and exported to display in QGIS. Spatial kriging of inorganic carbon was performed in R. The manually delineated meander polygons were converted to SpatialPixelsDataFrame using the sp package. A simple variogram model was fit to the natural log transformed inorganic carbon values with a spatial cutoff of 60 m. Kriging was then performed using the sample points, the meander SpatialPixelsDataFrame, and the fitted variogram model. The natural log transformed inorganic carbon values were then back transformed and the kriged map exported for visualization in QGIS.

## Data Availability

Representative genomes in the subspecies level set (98% ANI) can be accessed at https://ggkbase.berkeley.edu/ER15_ALL_curated_dRep98/organisms and representative genomes in the species level set (95% ANI) can be accessed at https://ggkbase.berkeley.edu/ER15ALL_ERML16_dRep95/organisms. Please note ggKbase is a ‘live’ site, genomes may be updated after this publication. Raw sequence reads for all metagenomes and metatranscriptomes included in this study can be accessed in the NCBI Bioproject Database using the umbrella accession number (PRJNA630765). Supplementary Table 1 includes NCBI Bioproject accession numbers for individual metagenomes metatranscriptomes.

## Acknowledgements

We are grateful to Chad Hobson, David McGrath, and Rosemary Carrol for support in the field during samples collection; and to Joel Rowland (Los Alamos National Lab) and Helen Malenda (USGS) for useful insights. We thank the Rocky Mountain Biological Laboratory for lab space at the field site. This work was supported as part of the Watershed Function Scientific Focus Area funded by the U.S. Department of Energy, Office of Science, Office of Biological and Environmental Research under Award Number DE-AC02-05CH11231. Sequencing was conducted at the Joint Genome Institute (a DOE Office of Science User Facility) under a CSP award.

