## Supplementary Information for "Meanders as a scaling motif for understanding of floodplain soil microbiome and biogeochemical potential at the watershed scale"

Current affiliation:
^ Department of Microbiology and Immunology, Stanford University, Palo Alto, CA, USA

File contents:
Supplementary figures 1-7
List of supplementary tables 1-14
List of supplementary data 1-8

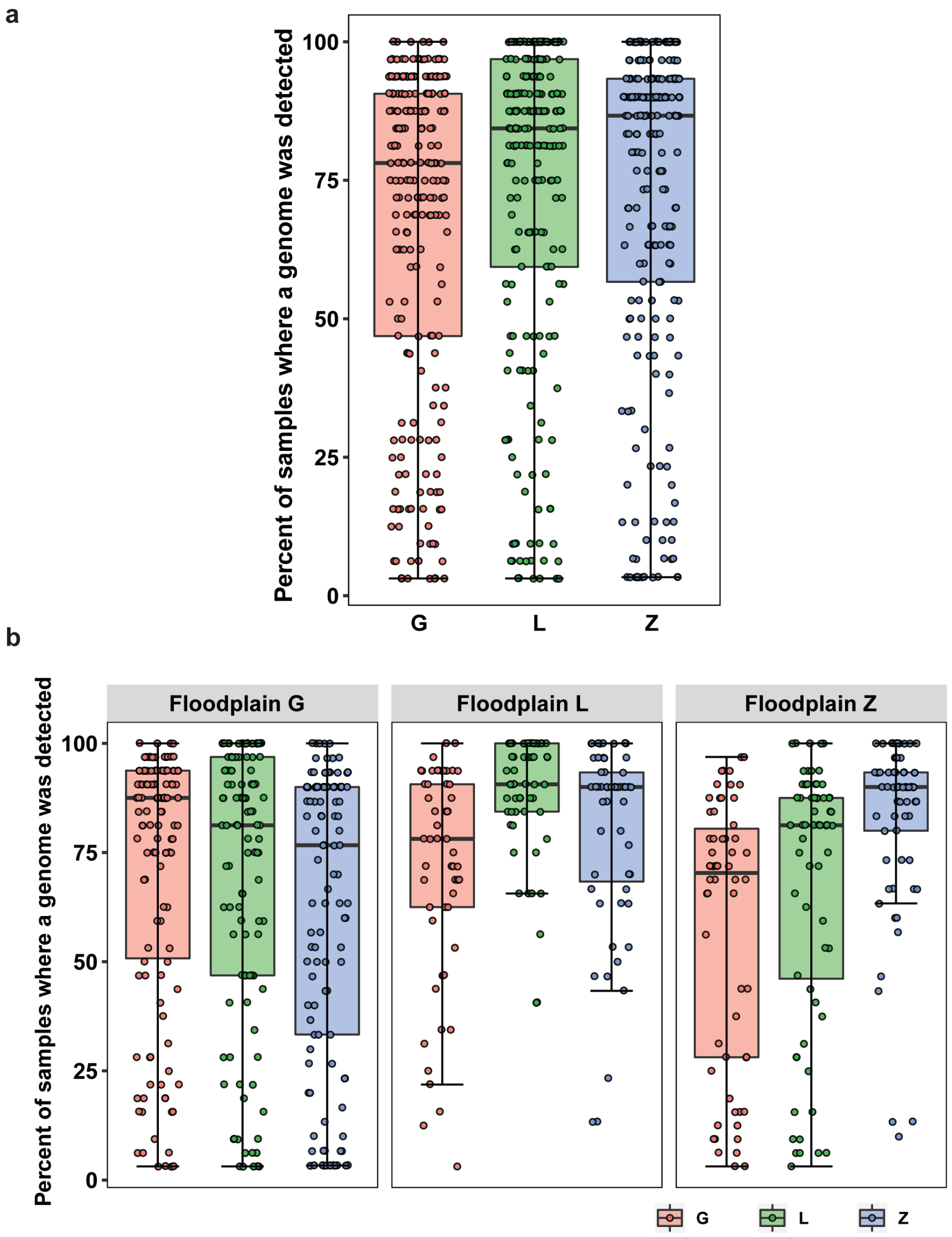


**Supplementary Figure 1.** Percent of samples within each floodplain where a genome was detected at the sub-species level (98% ANI). Presence or absence was determined based on Hellinger transformed abundance (average coverage ≥ 0.01). (**a**) Detection regardless of where the genome was reconstructed from. (**b**) Detection according to floodplain of origin. Genomes were detected in a higher number of samples from a given floodplain if they were reconstructed from a sample within that floodplain.

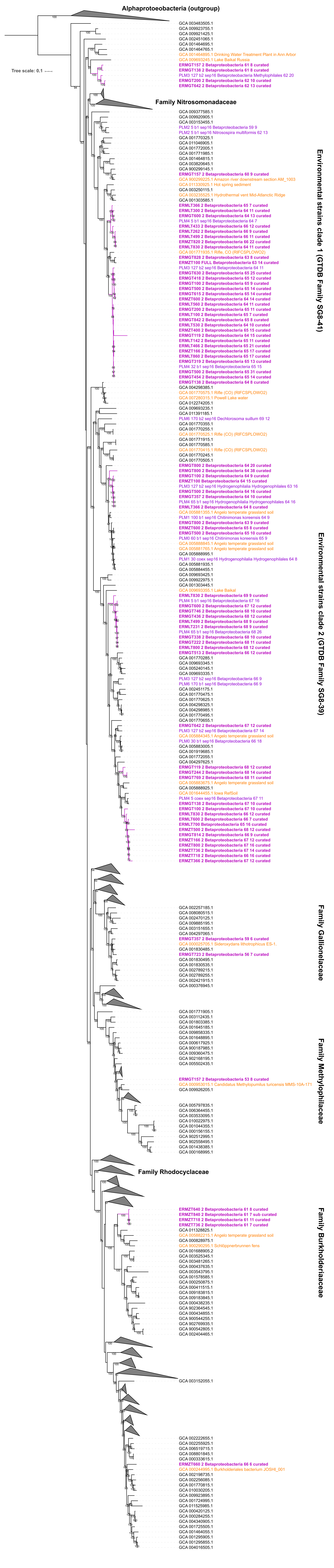
Alphaprotoeobacteria (outgroup)
Tree scale: 0.1
Family Nitrosomonadaceae
Environmental strains clade 1 (GTDB Family SG8-41)
Environmental strains clade 2 (GTDB Family SG8-39)
Family Gallionellaceae
Family Methylophilaceae
Family Rhodocyclaceae
Family Burkholderiaceae

**Supplementary Figure 2.** Concatenated ribosomal proteins IQ-TREE of Betaproteobacteria at the sub-species level (98% ANI) and ~ 1540 reference genomes from the NCBI. East River Betaproteobacteria are shown in bold magenta font (from this study) and violet font (from [1]). Some environmental sequences related to East River Betaproteobacteria are highlighted in orange, and next to the accession number is the environment of origin. Clades at the family level follow GTDB taxonomy [2] for additional reference.

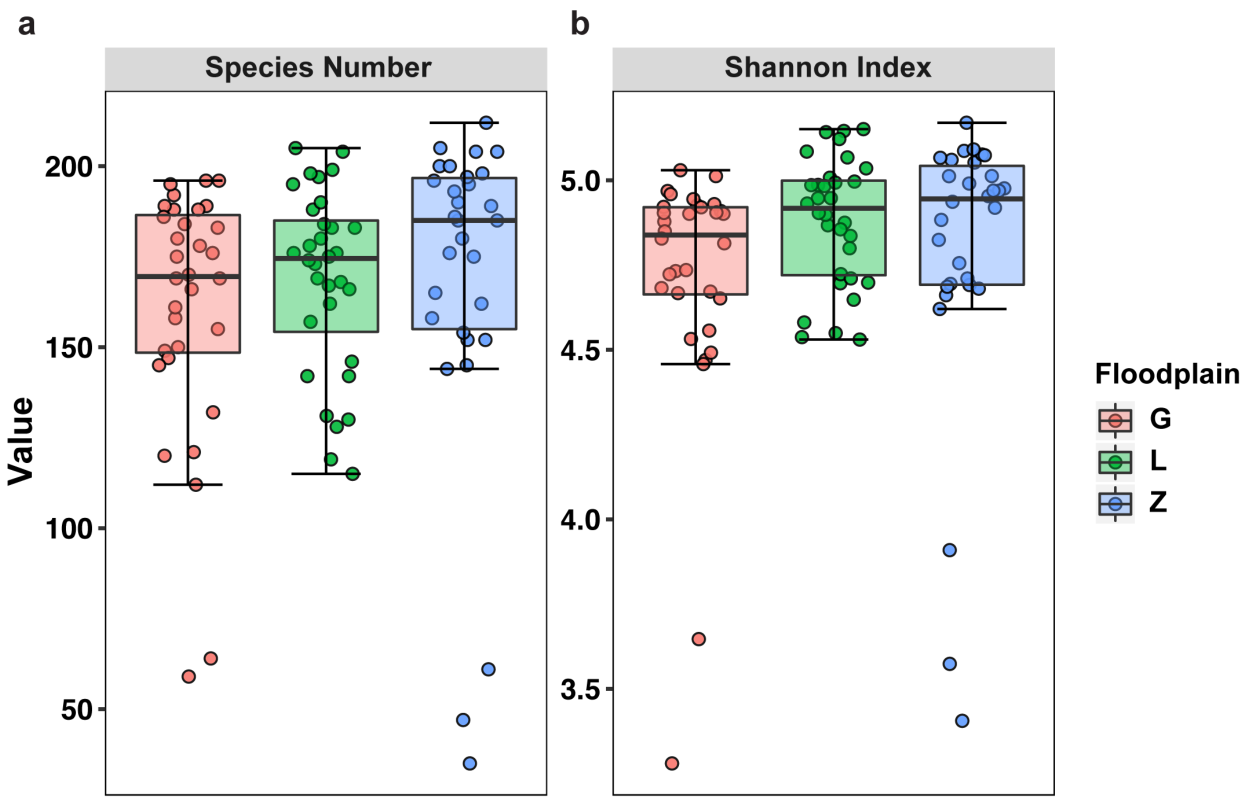


**Supplementary Figure 3.** Diversity indices calculated for a set of representative genomes at the sub-species level (98% ANI).

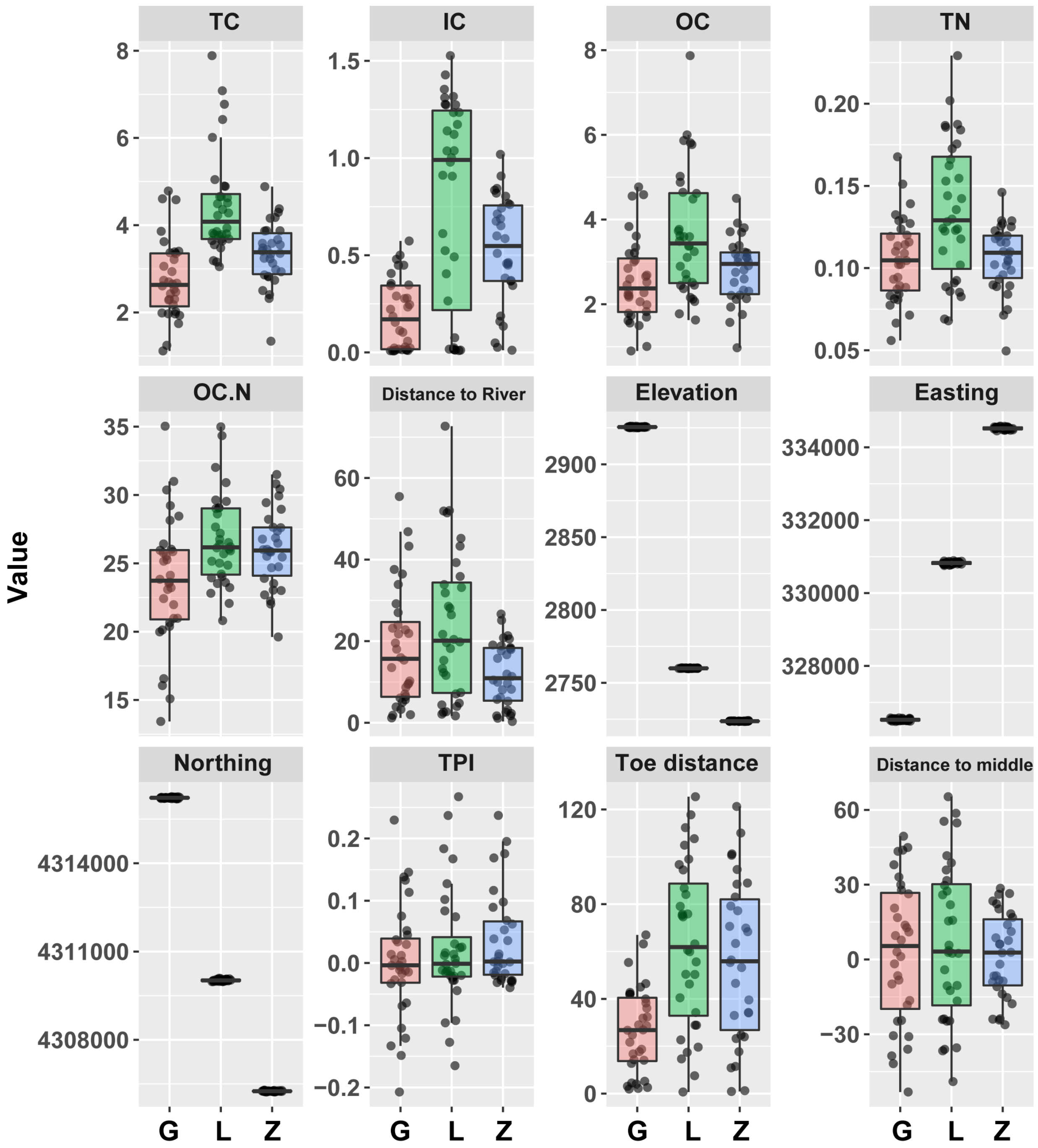


**Supplementary Figure 4.** Environmental variables used in fourth corner analysis: solid face chemistry from soil samples collected in 2015 including total carbon (TC; %), inorganic carbon (IC; %), organic carbon (OC; %), total nitrogen (TN; %), total carbon to total nitrogen ratio (OC:N) and measures associated with sample site locations: distance to river, elevation, easting, northing, topographic position index (TPI), and distance to the inner bank edge (or toe distance) and distance to middle of the meander-bound floodplain.

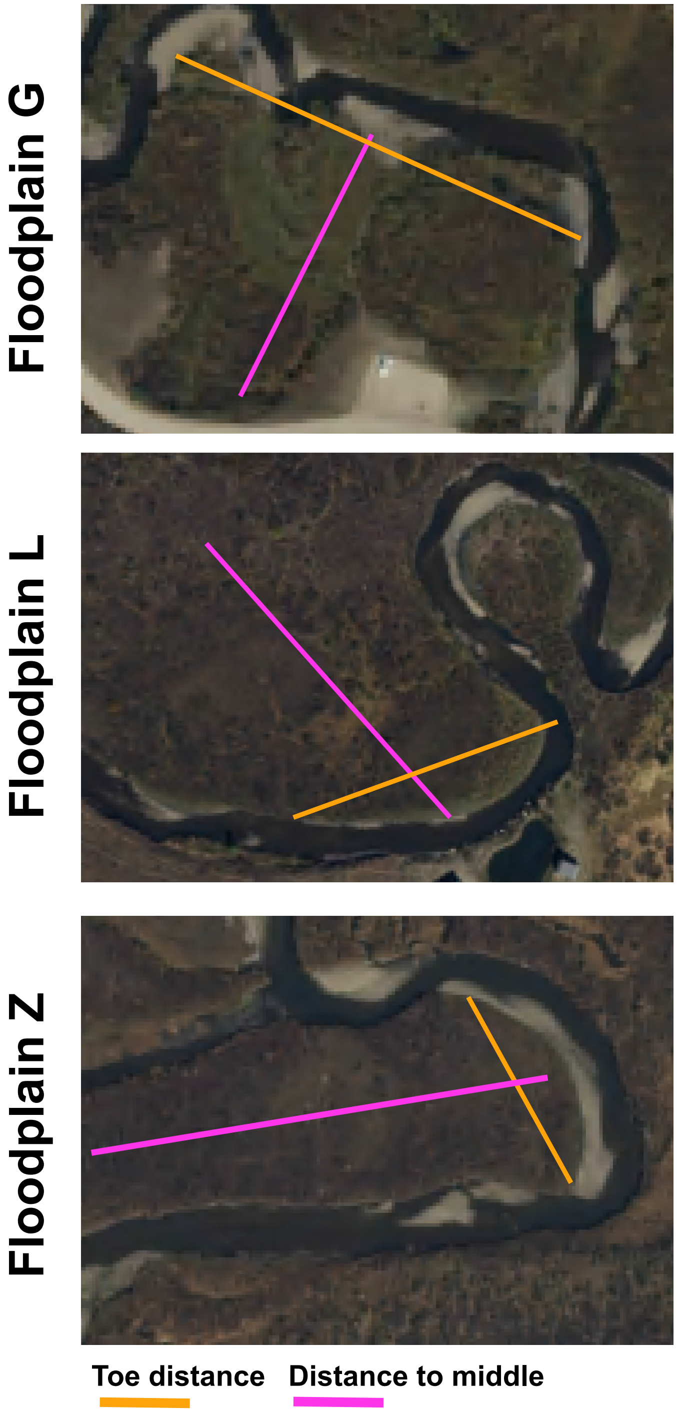


**Supplementary Figure 5.** Diagram representing imaginary lines used to determine distance to the middle of the meander-bound floodplain and distance to the inner bank edge (toe distance) as alternative measures of samples position on the floodplains.

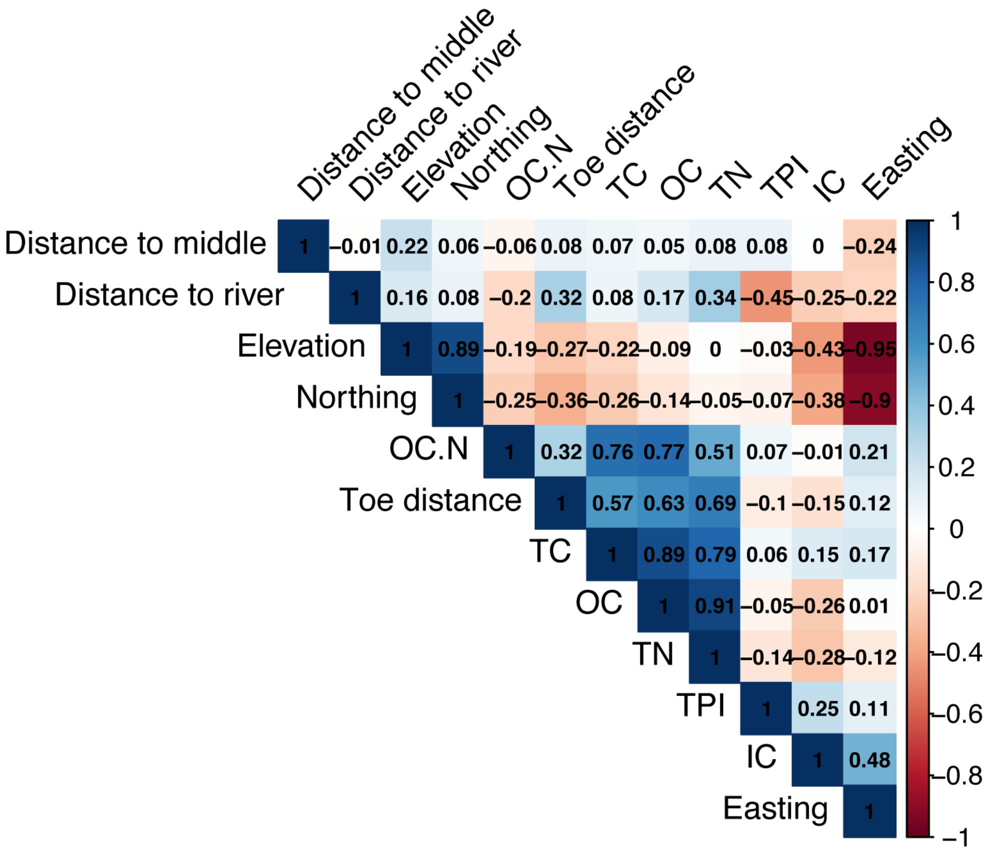


**Supplementary Figure 6.** Spearman's correlation among environmental variables (2015).

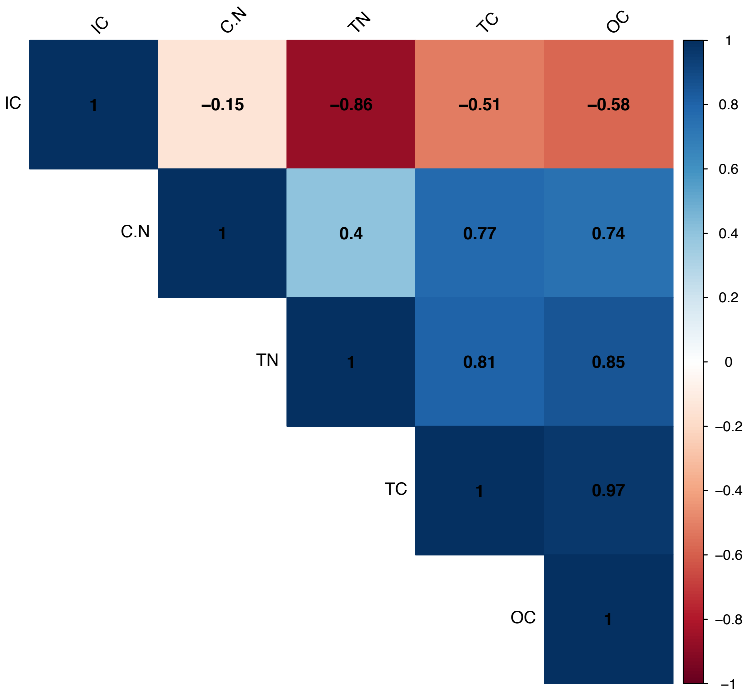


**Supplementary Figure 7.** Spearman's correlation among environmental variables (2016).

**List of Supplementary Tables** (separate files)

**Supplementary Table 1**: Sample sequencing and assembly information, and NCBI accession numbers.

**Supplementary Table 2**: Representative genomes (of 248 sub-species clusters at 98% ANI) information. This set of genomes was reconstructed from soil samples collected in September 2015.

**Supplementary Table 3**: Representative genomes (of 215 species clusters at 95% ANI) information. This set of genomes includes genomes reconstructed from samples collected in September 2015 and September 2016.

**Supplementary Table 4**: Presence/Absence of gene homologs identified among 248 representative genomes using selected HMMs. Includes rules used to define functions. Only "Present" considered for downstream analyses.

**Supplementary Table 5**: Results of the indicator species analysis.

**Supplementary Table 6**: Presence/Absence of functions among 248 representative genomes.

**Supplementary Table 7**: Solid phase chemistry for subset of samples collected in September 2016 with paired metatranscriptomes.

**Supplementary Table 8**: Carbohydrate active enzymes (CAZYs) in the glycosyl hydrolases (GH), carbohydrate esterases (CE), polysaccharide lyases (PL), and auxiliary activities (AA) classes detected among the 215 representative genomes (e- value cut-off 1e-20).

**Supplementary Table 9**: Unique sequences from both genome sets confirmed to be homologous to NiFe hydrogenases groups 1, 2 or 3 based on a phylogenetic analysis.

**Supplementary Table 10**: Unique sequences from both genome sets confirmed to be homologous to FeFe hydrogenases groups A, B or C based on a phylogenetic analysis. Sequences identified among the 248 representative genomes were used in analyses based on presence/absence. Sequences identified among the 215 representative genomes were used in transcript analyses.

**Supplementary Table 11**: Unique sequences from both genome sets confirmed to be homologous to FdhA/FdoG/FdhF/FdwA, NxrA, NapA, and NarG based on a phylogenetic analysis.

**Supplementary Table 12**: Unique sequences from both genome sets confirmed to be homologous to CoxL based on a phylogenetic analysis. Subtype is indicated next to the sequences.

**Supplementary Table 13**: Unique sequences from both genome sets confirmed to be homologous to methanol dehydrogenases.

**Supplementary Table 14**: Unique sequences from both genome sets homologous to DsrAB. Subtype is indicated next to the sequences.

**List of Supplementary Data** (separate files)

**Supplementary Data 1**: Concatenated ribosomal proteins phylogenetic tree of Betaproteobacteria among 248 representative genomes of sub-species level clusters at 98% ANI.

**Supplementary Data 2**: Concatenated ribosomal proteins phylogenetic tree of 215 representative genomes of species level clusters at 95% ANI.

**Supplementary Data 3**: Phylogenetic tree of NiFe hydrogenases groups 1, 2 and 3 reference sequences and homologs identified among unique sequences from both genome sets.

**Supplementary Data 4**: Phylogenetic tree of FeFe hydrogenases groups A, B and C reference sequences and homologs identified among unique sequences from both genome sets.

**Supplementary Data 5**: Phylogenetic tree of DMSOR superfamily reference sequences and homologs identified among unique sequences from both genome sets.

**Supplementary Data 6**: Phylogenetic tree of CoxL reference sequences and homologs identified among unique sequences from both genome sets.

**Supplementary Data 7**: Phylogenetic tree of PQQ-containing alcohol dehydrogenases reference sequences and homologs identified among unique sequences from both genome sets.

**Supplementary Data 8**: Phylogenetic tree of concatenated DsrAB reference sequences and homologs identified among unique sequences from both genome sets.
